# A theory of evolutionary dynamics on any complex spatial structure

**DOI:** 10.1101/2021.02.07.430151

**Authors:** Yang Ping Kuo, César Nombela Arrieta, Oana Carja

## Abstract

Understanding how the spatial arrangement of a population shapes its evolutionary dynamics has been of long-standing interest in population genetics. Most previous studies assume a small number of demes connected by migration corridors, symmetrical structures that most often act as well-mixed populations. Other studies use networks to model the more complex topologies of natural populations and to study the structures that suppress or amplify selection. However, they usually assume very small, regular networks, with strong constraints on the strength of selection considered. Here we build network generation algorithms, evolutionary simulations and derive general analytic approximations for probabilities of fixation in populations with complex spatial structure. By tuning network parameters and properties independent of each other, we systematically span across network families and show that both a network’s degree distribution, as well as its node mixing pattern shape the evolutionary dynamics of new mutations. We analytically write the relevant selective parameter, predictive of evolutionary dynamics, as a combination of network statistics. As one application, we use recent imaging datasets and build the cellular spatial networks of the stem cell niches of the bone marrow. Across a wide variety of parameters and regardless of the birth-death process used, we find these networks to be strong suppressors of selection, delaying mutation accumulation in this tissue. We also find that decreases in stem cell population size decrease the suppression strength of the tissue spatial structure, hinting at a potential diminishing spatial suppression in the bone marrow tissue as individuals age.

## Introduction

Novel microfuidics and organoid technologies (Simian and Bissell, 2017; Holloway et al., 2019) allow us to start building biological scaffolds that control the spatial topology of a molecular or cellular population. In order to make full use of these innovations, we need a rigorous theory of how the spatial structure of a population shapes its future evolutionary dynamics. This will allow us to design structures that either amplify the selective benefit and spread of beneficial mutations, or structures that suppress the spread of deleterious variants.

There is a large body of literature in population genetics theory studying the role of population structure in shaping evolutionary outcome, starting from the classic 1975 paper of Slatkin and Maruyama (Slatkin and Maruyama, 1975). However, most previous modeling approaches that incorporate spatial patterns of variation usually only assume a few demes (patches) and symmetrical structures (Wright, 1943; Kimura and Weiss, 1964; Carja et al., 2014), simple topologies that can be embedded into two-dimensional continuous Euclidean space. In most cases, these simple topologies do not change fixation probabilities and rates of evolution compared to well-mixed populations (Maruyama, 1970; Slatkin, 1981). These symmetrical structures fundamentally fail to capture the complex pattern of interaction and the variance in local selection pressure present in natural populations, as well as in emerging spatial cellular and molecular atlases (Regev et al., 2018; Snyder et al., 2019; Uhlen et al., 2010).

Studying more complex topologies, ones for which there exists no homeomorphism to the well-behaved two-dimensional Euclidian space, becomes a much harder mathematical problem. These topologies are best represented using networks and we can use the mathematical formalism of the Moran birth-death process on graphs (Lieberman et al., 2005; Carja and Creanza, 2019) to explore how spatially-structured patterns of interaction and replacement drive the composition of populations and shape the outcome of the evolutionary process. Graph theory has been successfully used to study patterns of spatial variation and interaction across a wide range of scientific fields, from the social sciences to brain science (Centola, 2010; Bassett and Bullmore, 2006; Sood and Redner, 2005). Network approaches have mostly been used in population genetic theory to study evolutionary game-theory, with nodes of the network representing individuals playing against each other and links representing the network of interaction between them (Nowak, 2006).

The power of these graph-theoretical approaches is hinted at in evolutionary studies of the Moran birth-death process on small, regular graphs. Here, a population of individuals is located on a graph and the links of a node cell indicate the neighboring nodes that can be replaced by its offspring (Lieberman et al., 2005). The spatial structure of the population is therefore represented through the graph structure and is studied in comparison to a well-mixed population. Initial studies (on very small, regular graphs) observe large differences between networks in the fixation probability of new mutants, compared to well-mixed populations (Lieberman et al., 2005; Hindersin and Traulsen, 2015), a marked departure from previous deme-based models. Some graphs are suppressors of selection, graphs that reduce the fixation probability of advantageous mutations, while increasing it for deleterious mutants (Lieberman et al., 2005; Hindersin and Traulsen, 2015; Hindersin et al., 2016a). Other graphs can be classified as amplifiers, increasing rates of evolution. One of the first general results, the isothermal theorem, states that in graphs where the propensity for change in each node is exactly the same, the fixation probabilities of new mutations are the same as in well-mixed populations. The assumptions of the isothermal theorem, however, sit on a knife edge; make small perturbations and the assumptions no longer hold (Lieberman et al., 2005).

While these initial studies hint at the promise of graph theoretical approaches, analytic results with predictive power have been very difficult to derive. Most prior results either rely on very small networks (where build and solve time scale exponentially with population size, making them unsuitable for the study of networks of more than 30 nodes (Hindersin et al., 2016a)) or invading mutants in the limit of neutrality, results that do not scale to generality (McAvoy and Allen, 2019). What are the probabilities of fixation for a new mutation as a function of where and when it appears in much larger, and more biologically-realistic spatial networks? We lack a predictive theoretical foundation of evolutionary outcome for populations that cannot simply be divided into a handful of separate patches, that is, populations with complex network-type spatial topologies.

Here, we develop a general theory of evolution on complex network structures and identify graph properties that control suppression or amplification of selection, either leading evolution to a stand-still or accelerating the evolutionary process. Linking network topology to evolutionary dynamics is complicated by the fact that networks differ in many structural properties. We build graph generation computational methods that allow the ability to systematically tune network statistics and study their role on probabilities and times to fixation for new mutations in the population, mathematical proxies for evolutionary outcome. Our algorithms combine simulated annealing procedures and degree preserving edge swapping (Taylor, 1981) to continuously tune network properties one at a time, while keeping other properties constant. This allows us to fully understand the role of specific network parameters, as well as translate the meaning of the relevant parameters for any given network, across different graph-families. We also derive general analytic approximations for probabilities of fixation on any graph structure, based on parameters describing graph degree distribution and mixing pattern (the probability that edges of different degrees are connected).

Using our simulations and analytical approximations, we find that knowing the degree distribution alone is surprisingly not enough to determine the fixation probabilities and times to fixation since graphs with the same degree distribution, but different mixing pattern can exhibit very different evolutionary outcome. Importantly, we analytically derive the relevant selective parameter, predictive of rates of evolution, for any given network without making restrictive assumptions on network type, size or selective advantage of the new invading variant.

In addition to the purely theoretical interest of the questions presented above, we also use our theoretical results to analyze rates of evolution in the stem cell populations of the bone marrow (Klein and Simons, 2011; Tomasetti et al., 2017). We use recent microscopy data sets (Coutu et al., 2018; Gomariz et al., 2018) to build the spatial stem networks of the bone marrow and we find that these networks are strong suppressors of selection, across a wide range of parameter choices and regardless of the type of the assumed birth-death process. Moreover, we find decreasing suppression with decreasing population size, hinting at a potential decrease in the suppressive properties of the spatial structure as individuals age. We also compare the evolutionary dynamics in these biological networks with the ones in online social networks and highlight the significant discrepancies that appear for the Birth-death process. While biological networks tend to suppress regardless of the order of birth and death in the Moran process, social networks exhibit very different dynamics depending on whether birth occurs before death or vice-versa.

There is a growing interest in the design of networks that optimize probabilities and times of spread of advantageous mutations in a population (for example in experimental evolution or for biotechnological applications), or the spread of cultural information (or misinformation) across social networks (Lieberman et al., 2005; Möller et al., 2019). Our results provide a general predictive theoretical framework for understanding the relevant network statistics for a population’s future evolutionary dynamics.

### Model

We use a Moran-type model to describe changes in allele frequencies in a finite population of constant size *N*. Each individual’s genotype is defined by a single biallelic locus *A/a*, which controls the individual’s reproductive fitness. An individual with the *A* allele is assumed to have fitness one, while an individual with allele *a* has assigned fitness (1 + *s*).

We use graphs and networks to represent the population structure. Every individual occupies a node in the graph, while the edges between two nodes represent the spatial or replacement structure of the population. One can also interpret the nodes as homogeneous subpopulations, fixed in space, with the edges representing corridors of migration and replacement between these subpopulations. Here, the time scale of a new beneficial mutation spreading and fixing in the node subpopulation is assumed to be much smaller than the time scale of mutational spread across the nodes.

Every generation, the population composition can change due to the interaction between selection and genetic drift and we update the allele frequencies using two different update scenarios (**Figure 1A**). In the first update scenario, we assume reproduction occurs before death (we denote this by the Birth-death *Bd* scenario). At every time step, we first select one node for reproduction, with probability proportional to fitness, from the entire population. We then randomly select one of its neighbors for death and vacate the node for the new offspring. In the second update scenario, denoted as the death-Birth *dB* update, we first select a node at random from the population to be vacated and then choose one of its neighbors for reproduction, with probability proportional to fitness. We can also interpret these processes as beneficial mutant spilling out of their subpopulations and invading neighboring nodes through the migration corridors. Note that selection happens only when choosing the node to reproduce. This means that, in the Birth-death update, the nodes compete globally, at the population level, while in the death-Birth update, the selection step is local, the competing nodes are only the neighbors of the node randomly chosen for death. Due to these differences in global versus local competition, the two update rules have been shown to lead to drastically different evolutionary dynamics (Lieberman et al., 2005; Hindersin and Traulsen, 2015).

**Figure 1:**
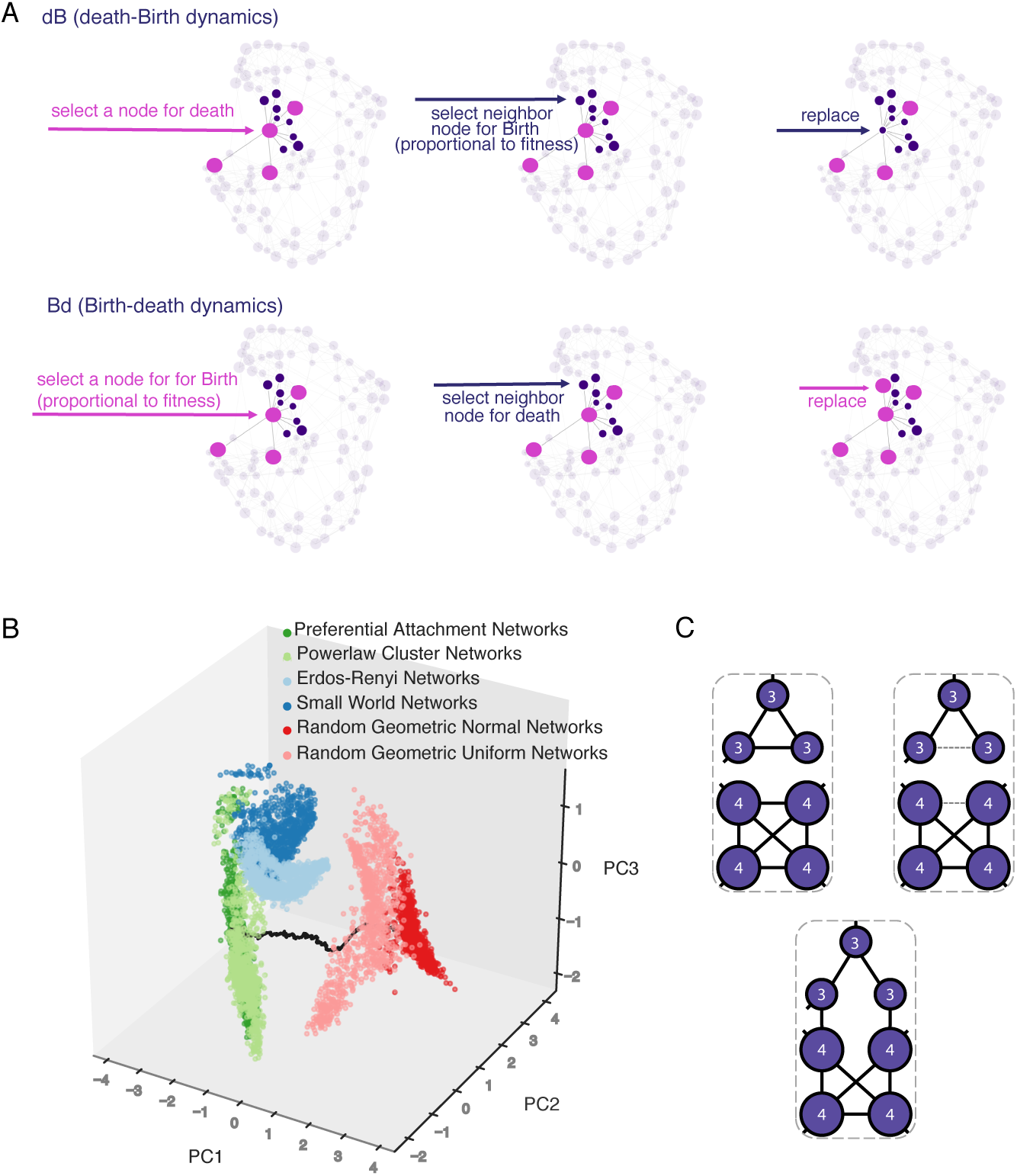
Illustration of the population birth death process update rules and graph rewiring methods. **Panel A** illustrates both the dB (death-Birth) and the Bd (Birth-death) update rules. In **Panel B**, we use principle component analysis on 6 graph characteristics (mean, variance, third moment, modularity, average clustering, and assortativity) to highlight the network families studied, as well as the trajectories between them. Each graph family shows clustering using the first three principle components (that explain 89% of the variance in PC space). The black line represents the trajectory in PC space as we rewire graphs starting from preferential attachment (PA) graphs, through powerlaw cluster networks (PLC) and uniform random geometric graphs, to normal random geometric graphs (RGG). **Panel C** illustrates the edge swap operation used to tune graph characteristics. At first, there are no edges connecting nodes of degree 3 to nodes of degree 4. Two edges are randomly selected to be disconnected and nodes that were “parallel” with respect to the two disconnected edges are then connected, thus preserving the number of edges. After the rewiring step, there are two edges connecting nodes of degree 3 to nodes of degree 4, and there is no longer a 4-clique. The degrees of the nodes, however, are preserved.

The graph structure therefore becomes a mathematical proxy for the population spatial topology or the population structure of interaction: nodes reproduce locally and the edge pattern imposes limits to variant spread. The graphs we consider here are unweighted and undirected. Initially, we assume the population fixed on the wild-type *A* allele. We introduce one mutant *a* individual at a random node at time *t* = 0. We ignore subsequent mutation, and therefore the population will eventually reach a monomorphic state where individuals of the same *A/a* allele occupy all nodes in the graph. The quantities of interest become the probability of fixation and the time to fixation of the invading allele *a*, informing on the rate of evolution of the population for a given graph topology.

Our goal here is to systematically study the role of the network structure in shaping rates of evolution by directly comparing these probabilities of fixation with the equivalent probabilities in a well-mixed population. To identify the relevant graph properties that either speed up or suppress adaptation, we characterize graphs through the lens of their main two components: nodes and edges. We can therefore think of graph properties as either node- or edge-centric. The main property of a node is the node degree (the number of neighbors the node is linked to) and the node degree distribution becomes an important global network property (Newman, 2003). Graph edges, on the other hand, can be categorized based on the type of nodes they connect and how often they connect nodes of different degrees. The mixing pattern of a graph (also called graph assortativity) is a global edge-centric graph descriptor that informs on the frequencies of each edge type (Newman, 2002). We use random graph generators and construct graphs that differ in their degree distributions and mixing patterns and thus span the known graph families (Barabási and Albert, 1999; Holme and Kim, 2002; Watts and Strogatz, 1998; Waxman, 1988; Penrose et al., 2003; Erdős and Rényi, 1960; Masuda et al., 2005). This ensures that our results are generalizable across graph families and graph properties (**Figure 1B**). Two graph generator families we will highlight are the Barabasi-Albert model of preferential attachment and the generalized random geometric model.

In preferential attachment (PA) graphs, each network starts with a single node and nodes are added sequentially until the population reaches size *N*. Each new node is added to the network and connected to other individuals with a probability proportional to the individual’s current degree to the power of a given parameter *β*. By adjusting the number of edges added each step (*m*) and the power of preferential attachment (*β*), this family of graphs allows for straightforward independent tuning of the moments of the degree distribution. Parameter *m* is the only parameter of the model that controls the first moment of the degree distribution. Parameter *β* controls the shape of the distribution, with the distribution being exponential when *β* = 0, stretched exponential when 0 *< β <* 1, and power law when *β* = 1 (Krapivsky et al., 2000). When *β* = 1, PA graphs exhibit the scale-free property (Barabási and Albert, 1999) and PA graphs are often used as a model to study the spread of information or cultural norms (Creanza et al., 2017).

To model a spatially structured population, we use generalized random geometric graphs (Waxman, 1988; Penrose et al., 2003). In this family of graphs, nodes have spatial positions randomly drawn from a probability distribution to model spatially homogeneous populations (using the uniform distribution) or populations with heterogeneous spatial density (using the normal distribution). Once the spatial locations of the nodes are determined, the generating algorithm iterates through all pairs of nodes. An edge is created between two nodes using a probability distribution based on pair-wise distance. Here we use an exponential distribution (the resulting graphs are known as Waxman graphs) and a heavy-side function where we connect two nodes if the distance is below a predefined threshold (denoted as random geometric graphs).

Existing graph generating methods do not allow for simultaneous tuning of the degree distribution and the node degree mixing pattern. For example, in PA graphs, one can smoothly change the shape of the degree distribution by changing the power of preferential attachment *β*. However, if we change this preference of connection to high degree nodes, we inevitably also change the graph’s mixing pattern and cannot independently study its role in amplifying or suppressing selection.

To allow for tuning of single parameters of the degree distribution as well as tuning of mixing patterns, independent of the degree distribution, we implement a sampling network generation algorithm based on simulated annealing (**Figure 1C**). The algorithm relies on a degree swapping operation on graphs (Taylor, 1981) and runs for a preset number of time steps. The algorithm can take any graph as input. The degree distribution of the graph is preserved, while other properties such as the mixing patterns of nodes are changed. Given a degree distribution, we want to make sure we explore a wide range of graph mixing patterns. To this end, we use degree Pearson correlation *r* to measure the mixing pattern in the graph. To find the graph that maximizes the degree correlation, given a fixed degree distribution, we accept an edge swap according to the criterion

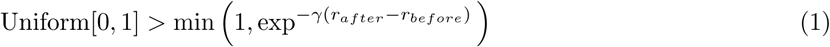

and reject the step otherwise. Here, 1*/γ* is the annealing temperature that controls how stringent the criterion must be and is decreased as the simulation proceeds. Intermediate graphs are periodically saved and we use the heuristic outlined in Gkantsidis et al. (2003) to periodically check that the graph is fully connected. This algorithm yields a set of graphs spanning a wide range of possible degree correlations, for a given graph degree distribution, thus allowing us to study the effects of mixing pattern on evolutionary dynamics.

Once the network structure is set, we use ensembles of at least 10, 000 Monte Carlo simulations, as well as analytic approaches as described in the next section, to compute the probability and times to fixation of the new allele *a*.

## Results

We study the probability of fixation of a new invader mutation *a* with fitness (1 + *s*) that appears in a random initial node of the network and compare it with the equivalent probabilities of fixation in well-mixed populations. We start by obtaining analytic approximations for the dB (death-Birth) update model and then discuss the important differences specific for the Bd (Birth-death) update rule. Intuitively, the dB dynamics applies when the evolutionary update is driven by available space being freed up, followed by local competition among the neighbors, whereas the Bd dynamics applies in cases where competition happens globally, but replacement is driven by the local pattern of interaction.

We begin by presenting the main ideas of our analytic approximation for the probability of fixation of the *a* allele. For the complete mathematical treatment, please see the **Supplementary Material**. Previous analytic approaches have either made use of the adjacency matrix of the network (which uniquely identifies the graph) and its associated transition probabilities (Hindersin et al., 2016a) or assumed that the evolutionary dynamics are in the limit of neutrality and a vanishing selection coefficient *s* (if weak selection is assumed, the probability of fixation can be approximated by treating it as the linear perturbation to the continuous coalescence, the dual of the Moran process under neutrality) (McAvoy and Allen, 2019; Allen et al., 2017). The former approach can provide closed form solutions for the fixation probability of *a*, but becomes intractable for large networks since it tracks a Moran process with 2^*N*^ states and the algorithm build and solve time both grow exponentially with population size (even for *N* = 23 nodes it can take several minutes (Hindersin et al., 2016a)). The latter approach reduces the problem from exponential to polynomial complexity in population size *N* (McAvoy and Allen, 2019; Allen et al., 2017), *however it performs poorly as we move away from the neutrality limit for the a* allele (**Supplementary Figure S1** and **S2**).

The approach we take here is to use the node degree distribution, and only keep track of the mutant frequencies *x*_*i*_ at all *N*_*i*_ nodes of the same degree *d*_*i*_. Let *D* = {*d*_1_, *d*_2_, …, *d*_*i*_, …} represent the set of all possible node degrees. While the degree distribution might not uniquely represent the network and some of the graph information is lost, this approach nonetheless greatly reduces the number of possible states in the Moran model (Ohtsuki et al., 2007; Sood and Redner, 2005). We denote the frequency of nodes of degree *d*_*i*_ in the population by *p*_*i*_. To model node degree mixing, we use *p*_*ij*_ to denote the probability that a node of degree *d*_*i*_ is connected to a node of degree *d*_*j*_. The probability of fixation of allele *a* is then approximated using the diffusion approximation, as described below (Kimura, 1962; Crow et al., 1970).

At every time point, *x*_*i*_, the frequency of the mutant at nodes of degree *d*_*i*_, increases by 1*/N*_*i*_ with probability 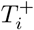 and decreases by 1*/N*_*i*_ with probability 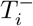. We can write

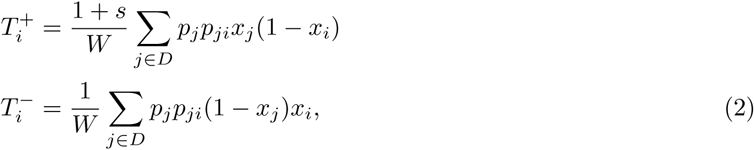

where *W* is the mean fitness of the individuals in the population.

We use these transition probabilities to find the mean and covariance of the change in *x*_*i*_ per unit time and use the backward Kolmogorov equation (Crow et al., 1970) to find the probability of fixation of the *a* allele for any initial mutant frequency 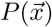:

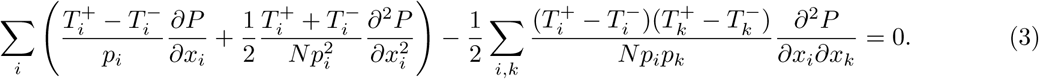

Here, the coefficient for the linear differential operator is quadratic in *x*_*i*_ and the coefficient for the quadratic differential operator is quartic in *x*_*i*_.

By using singular perturbation to linearize the coefficients of the differential equation (Gavrilets and Gibson, 2002), the solution to the partial differential equation in (3) for the Birth-death update model can be approximated by:

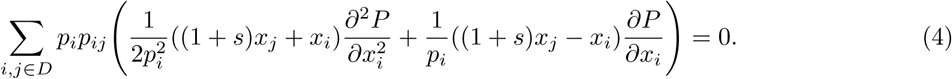

The death-Birth process shares a similar equation, given by

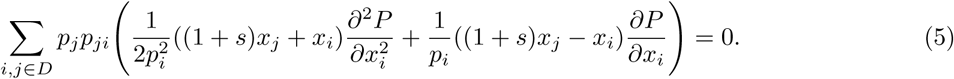

The only difference between the two equations is the change from *p*_*i*_*p*_*ij*_ to *p*_*j*_*p*_*ji*_. The solution for the Bd process can then be written as

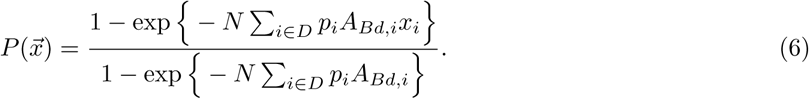

We can compute *A*_*Bd*_ by solving the following system of quadratic equations

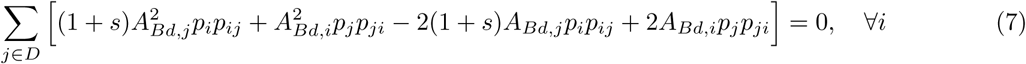

while for the dB update processes we need to solve

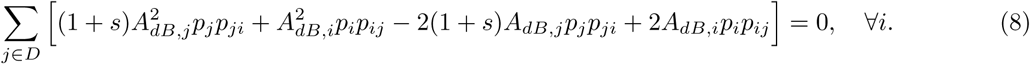

In the death-Birth process, the contribution to the fixation probability due to the degree distribution is on the order of the selection coefficient *s*, while the contribution due to degree mixing is on the order of *s*^2^. Therefore, knowing the degree distribution of the graph gives a good approximation to the probability of fixation, for weak *s*. Assuming 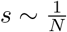, the probability of fixation can be approximated as

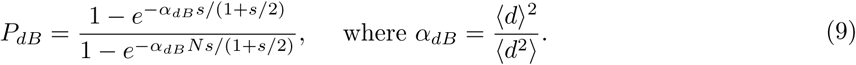

Here, ⟨*d*⟩ = Σ*p*_*i*_*d*_*i*_ and 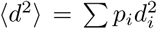 are the first and second moment of the degree distribution. This selection suppression or amplification factor *α*_*dB*_ can be used to measure how much the probability of fixation differs from that of well-mixed populations. If *α* = 1, the probability of fixation is identical to well-mixed populations. Graphs with *α >* 1 are amplifiers and *α <* 1 are suppressors. Our approximation shows that, for sufficiently weak selection, *α*_*dB*_, the suppression parameter for dB processes, is a function of the first and second moments of the degree distribution alone.

Solving equations (7) and (8), we show the accuracy of the analytic approximation in **Figure 2** using preferential attachment PA graphs. As the mean of the degree distribution increases, probability and time of fixation increase for the death-Birth process (and decrease for the Birth-death process) towards the well-mixed population limit. This makes intuitive sense: as the mean degree increases, the graph structure approaches that of a well-mixed population. In contrast, as the variance of the degree distribution increases, while keeping the mean constant, probabilities and times to fixation decrease monotonically for the dB process, and increase for the Bd process. The variance measures how heterogeneous the nodes are. At variance zero, all the nodes in the graph have the same number of neighbors, which means the graph is isothermal and the fixation probability is the same as that of well-mixed populations.

**Figure 2:**
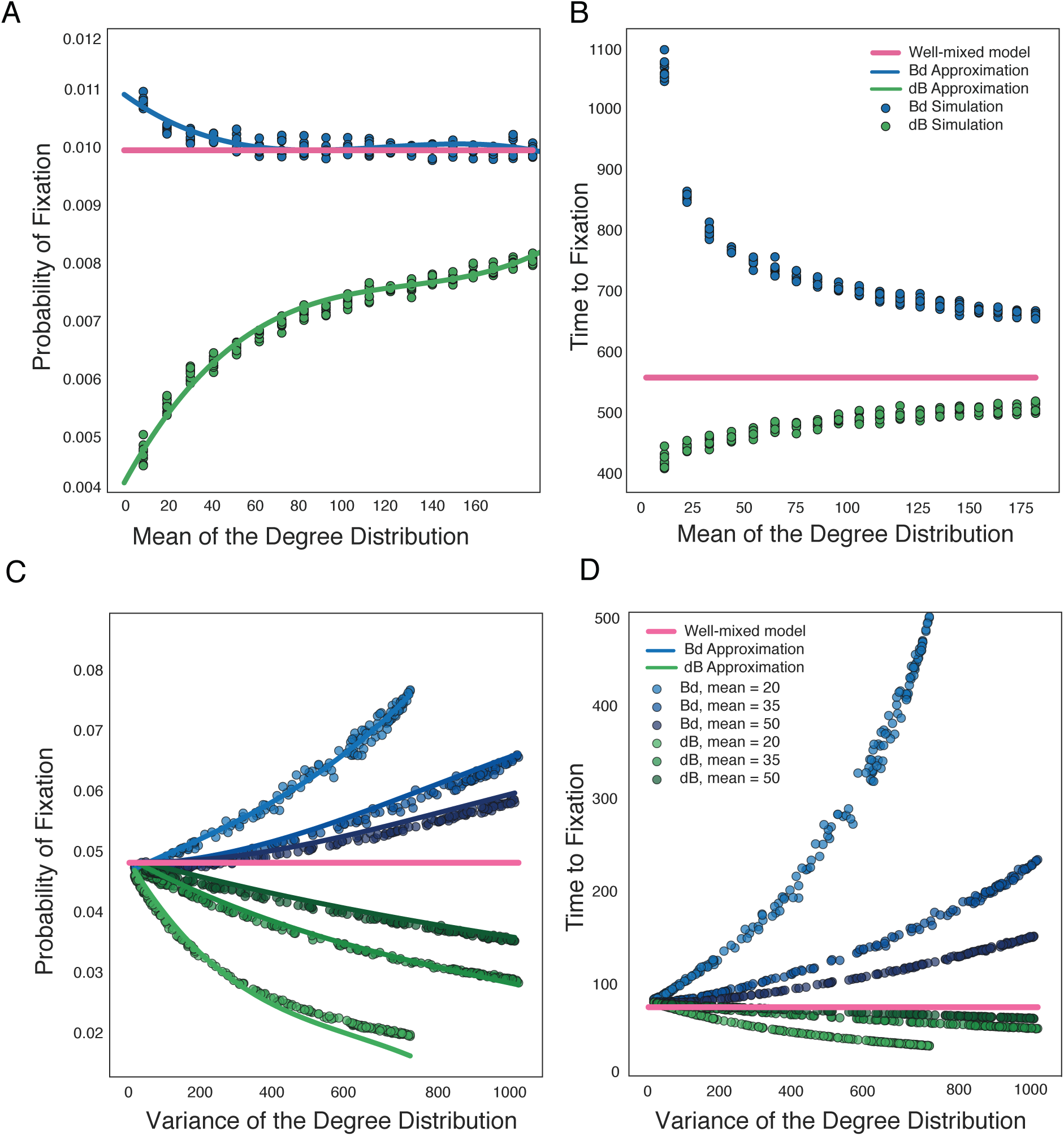
Role of the first moments of the degree distribution on evolutionary dynamics. The dots represent represent ensemble averages across 10^6^ replicate Monte Carlo simulations, while the lines represent our analytical approximations. **Panels A** and **C**: We plot the probabilities and times to fixation as a function of the mean of the degree distribution. We use preferential attachment PA graphs, graph size *N* = 1000 and *Ns* = 10. **Panels C** and **D**: We plot the probabilities and times to fixation as a function of the variance of the degree distribution, while keeping mean degree constant, as in the legend. Here, graphs are PA graps, *N* = 100 and *Ns* = 5.

Using the approximation in (9) for the death-Birth dB process, we show the probability of fixation of a new mutation *a* across multiple graph families in **Figure 3A**. As the effective selection parameter *α* increases, the fixation probability increases, reaching and crossing the well-mixed line when *α* equals to one. Intuitively, *α* quantifies the interplay between the mean and variance of the degree distribution, between how well-connected the nodes are and the network node heterogeneity.

**Figure 3:**
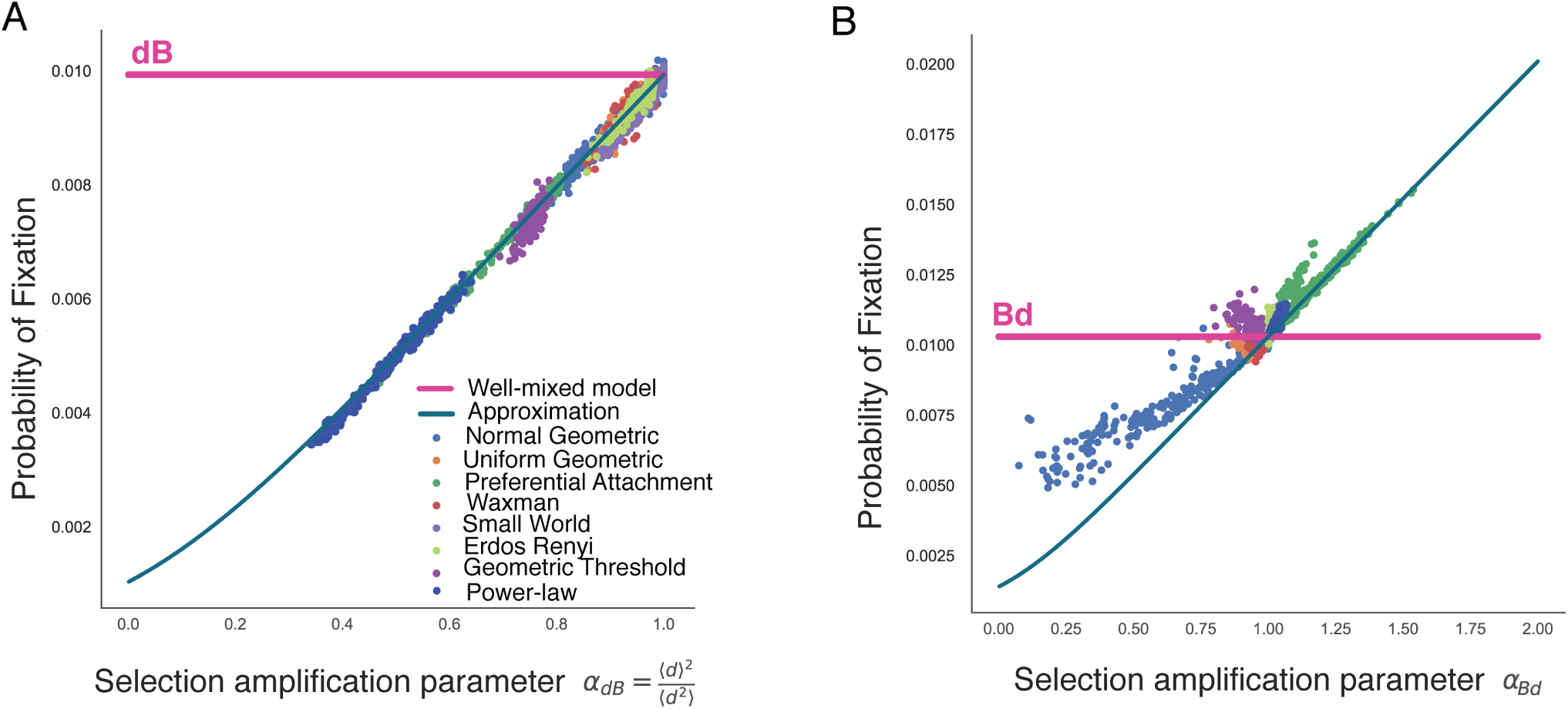
Probabilities of fixation across graph families. The fixation probability is shown on the y-axis as we vary the evolutionary quantity *α* on the x-axis. The dots represent represent ensemble averages across 10^6^ replicate Monte Carlo simulations, while the lines represent our analytical approximations. Here, graph size *N* = 1000 and *Ns* = 10. Each dot represents simulations on a distinct graph. The various colors represent graphs generated using different generation algorithms. **Panel A** shows results for the death-Birth process. **Panel B** shows results for the Birth-death process.

### Evolutionary role of the graph mixing pattern

For the Birth-death process, unlike the case of the death-Birth process where the effects of mixing pattern can be ignored under weak selection, network degree distribution and mixing pattern both contribute to the new mutation’s fixation probability. Similar to the death-Birth process, the contribution to the fixation probability due to degree distribution is again on the order of the selection coefficient of the new mutation *s*. However, in contrast to the death-Birth process, the graph mixing pattern has the same order of magnitude contribution as the graph degree distribution. Under selection 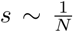, the probability of fixation can be approximated as

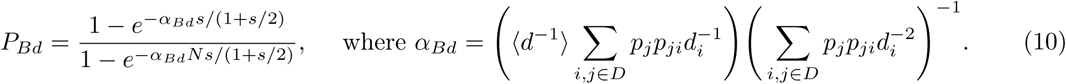

Here, 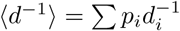 is the first inverse moment of the degree distribution.

For the Birth-death process, the *α*_*Bd*_ selection factor can be written as a function of parameters of the network wiring pattern and properties of its degree distribution. This approximation is shown in **Figure 3B**, alongside the results of Monte Carlo simulations. The fixation probability increases as the selection parameter *α* increases, with lower values for random geometric graphs and higher selection amplification for the preferential attachment graph family.

To understand the underlying network properties controlling evolutionary dynamics of new mutations, we need an intuitive understanding of the amplification factor in equation (10). The inverse moment ⟨*d*^−1^⟩ quantifies the shape of the degree distribution, while the rest of the parameters in equation (10) can be thought of as parameters that measure the graph’s assortativity or mixing pattern. A graph is assortative when a node of degree *d*_*i*_ preferentially attaches to other nodes of a degree similar to *d*_*i*_. A graph is called disassortative when the number of edges that connects nodes of degree *d*_*i*_ and nodes of dissimilar degree is higher than the expected number in randomly mixing graphs (Newman, 2002). Consider an edge swapping operation on a graph that breaks two edges: one between two nodes of degree *d*_*i*_ and one between two nodes of degree *d*_*j*_. Two edges that connect node of degree *d*_*i*_ and degree *d*_*j*_ are formed from the stubs. If *d*_*i*_ and *d*_*j*_ are dissimilar, such a rewiring step reduces the graph’s assortativity. Assuming the population size is large, the change in the *α*_*Bd*_ amplification factor can be written as

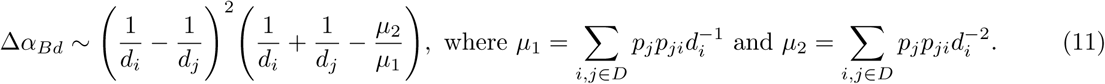

The magnitude of the change depends on the difference between the reciprocals of the degrees. If new edges are created between nodes of very dissimilar degrees, the change in the fixation probability can be significant. Since the change depends on the reciprocal, nodes of low degree have a disproportional effect on the change in amplification. The upper bound of *µ*_2_*/µ*_1_ is 1*/d*_*min*_, where *d*_*min*_ is the smallest degree of the graph. This means that if either *d*_*i*_ or *d*_*j*_ is close to the lowest degree, *α*_*Bd*_ is guaranteed to increase. In other words, the probability of fixation increases when there are more edges connecting nodes of low degrees to nodes of high degrees (disassortative graphs). An example of this is the star network, the strongest known amplifiers for undirected graphs (Möller et al., 2019). A star graph consists of a few nodes forming the center, while the rest of the nodes connect to the center nodes and form the vertices of the star. As a consequence, the nodes at the center have high degrees, while the rest tend to have significantly smaller degree, and the only edge type in the graph is between nodes of very different degrees. This type of graphs have the highest disassortativity, and strongest amplification of selection.

We can use this intuition to also explain the relative location of graph families in **Figure 3B**. Preferential attachment networks tend to be graphs with low assortativity (high disassortativity), with many hub-and-spoke structures (lower degree nodes connected to high degree nodes) and thus strong amplifiers. In contrast, normal geometric graphs with non-uniform spatial density tend to have high assortativity and thus tend to suppress the force of selection. This is due to the fact that nodes in high-density areas tend to be closer to each other and, assuming the density function is relatively smooth, neighbors tend to have similar degrees. Similarly, nodes in low spatial density areas tend to have few neighbors (low degrees), and so do their neighbors. Therefore, in spatial graphs with nonuniform spatial density, nodes are connected with neighbors of similar degree. This explains the suppression effect of this network family.

Knowing the degree distribution alone is not enough to determine the fixation probability since graphs with the same degree distribution but different mixing pattern exhibits different evolutionary dynamics. To illustrate the effects of assortativity and mixing pattern on fixation probabilities without the influence of degree distribution and graph generating method, we use edge swap operations to sample graphs with different mixing patterns, while keeping the degree distribution the same. We use graphs generated from different generating methods as input graphs to ensure generalization across graph families. For the same degree distribution, the spread of values for the fixation probability due to the effect of degree mixing can be substantial (**Figure 4A**). Here, all dots of the same color represent graphs from the same starting graph family and are altered by the edge swap sampling method with different end mixing patterns. We use the variance of the degree distribution as a measure for the shape of the distribution. Although the mean degree is not shown, dots of the same color share the same mean.

**Figure 4:**
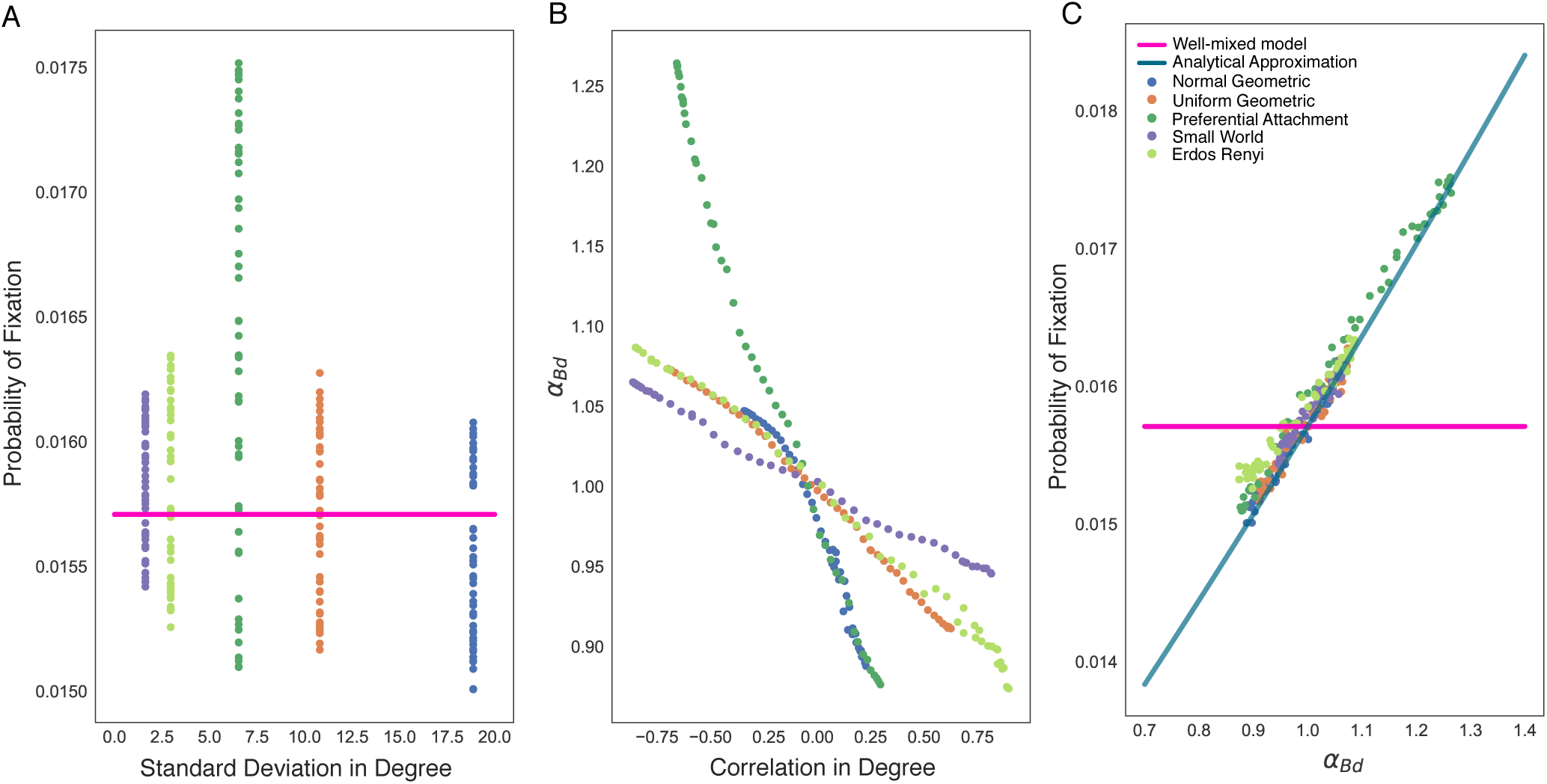
Effects of the mixing pattern on the Bd probability of fixation. The dots represent represent ensemble averages across 10^6^ replicate Monte Carlo simulations, while the lines represent our analytical approximations. Here the degree distribution is held constant as we vary the mixing pattern of the graphs, *N* = 100 and *s* = 0.1. Color indicate the graph family (with the same degree distribution) as in the legend. Mixing pattern of the graph is tuned using edge swapping operations. **Panel A**: Given a fixed graph degree distribution, there is a large range for the fixation probability of a new mutation on the network due to differences in graph mixing pattern. **Panel B**: The selection parameter *α*_*Bd*_ we derive captures these differences in graph mixing pattern and degree correlation. **Panel C**: The equivalent of **Figure 3B**, where we keep the degree distribution constant and just vary the mixing pattern for graphs spanning the illustrated graph families.

We use degree Pearson correlation *r* as a measure of the mixing pattern in the graph. We maximize and minimize the degree correlation to obtain an ensemble of graphs with the same degree distribution but different mixing pattern. When *r* = 1, the network has perfect assortative mixing patterns, while *r* = −1 corresponds to the case of a disassortative network. The contribution of the mixing pattern in the amplification constant in equation (10) is also a measure of assortativity (**Figure 4B**). Here *r* and *α*_*Bd*_ have a negative correlation, as expected. For graphs with the same degree distribution, the graphs that have low assortativity (high *α*_*Bd*_) have a higher probability of fixation (**Figure 4C**). The difference between **Figure 4C** and **Figure 3B** is that in **4C** we keep the degree distribution constant. Therefore, both node types and edge types in a graph contribute to evolutionary dynamics on the graph structure.

### Increased suppression of selection in large populations

While it has been previously claimed that under the Birth-death process most random graphs are amplifiers of selection (Hindersin and Traulsen, 2015), our results show that a large fraction of Birth-death graphs are suppressors of selection. The discrepancy in the results is due to the different population sizes considered. In this section, we study the effects of population size on fixation probabilities using two types of graphs: star graphs, known to be the strongest undirected amplifiers, and detour graphs, strong suppresors (Möller et al., 2019).

A detour graph consists of a completely connected central cluster and a cycle part (see **Figure 5**). These graphs have a low probability of fixation due to their high assortativity, since the graphs only have two edges connecting nodes of different degrees. We show that the fixation probability depends on the size of the central cluster, i.e. the length of the detour. To find the cluster size that minimizes the probability of fixation, we use the solution to the diffusion equation (47) in the **Supplementary Material**, derived using regular perturbation (Zhivotovsky and Feldman, 1993):

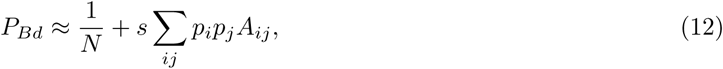

where *A*_*ij*_ satisfy the following system of linear equations

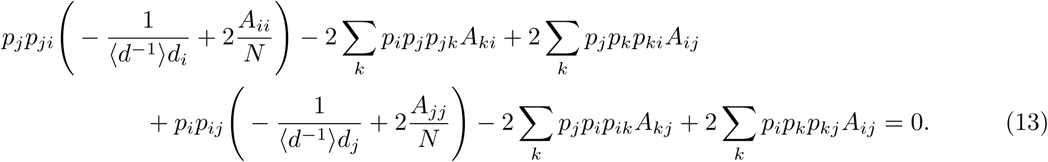

The size of this system of equations is |*D*|(|*D*| + 1)*/*2, where |D| is the number of unique degrees in the graph. Since a detour graph has only two types of degrees, we only need to solve a system of 3 equations for *A*_11_, *A*_12_, and *A*_22_. The only variable that influences the probability of fixation in the detour graphs is the length of the detour. We plot the difference in probabilities of fixation for detour graphs of different sizes and the well-mixed population against the length of the detour in **Figure 5A**. When detour length equals zero, we have the complete graph where the difference in the probability of fixation is zero. Since detour length is maximized in a ring graph, the probability of fixation initially decreases as the length of the detour is increased, reaching a minimum, before increasing towards the well-mixed control. It can also be observed that the minimum decreases with population size. The regular perturbation approximation is used instead of equation (10) since, while the approximation predicts the magnitude of suppression on detour graphs, the minima are shifted slightly to the left towards well-mixed. Mathematically, this is due to the fact that singular perturbation deviates from the solution of the diffusion equation when the exchange of individuals between two sub-populations is weak (Whitlock and Gomulkiewicz, 2005).

**Figure 5:**
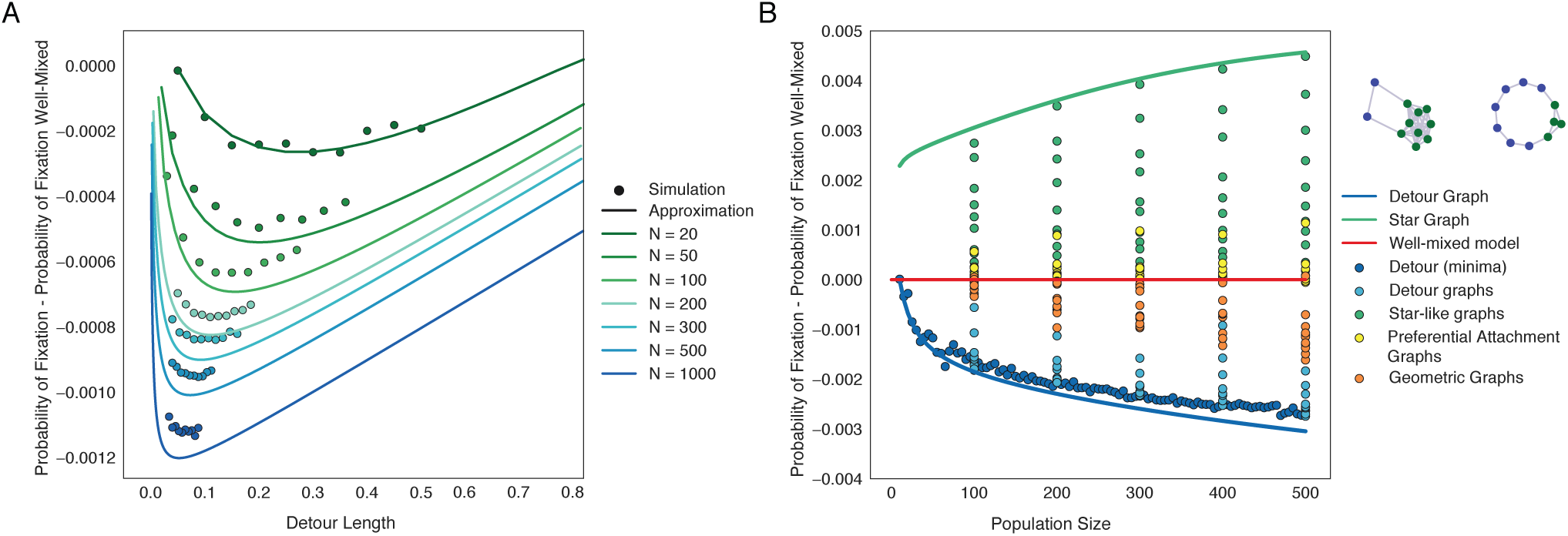
Detour and star graphs are graph families that minimize and maximize of probability of fixation under under the Bd process. The dots represent represent ensemble averages across 10^6^ replicate Monte Carlo simulations, while the lines represent our analytical approximations. Detour graphs are the strongest suppressors under weak selection. Detour graph consists of a completely connected central cluster and a cycle part (see the two illustrated networks). In **Panel A**, we plot the probability of fixation on detour graphs with varying lengths of the detour cycle. Here *s* = 0.002. In **Panel B**, we plot the probability of fixation across graph types, with respect to population size, to illustrate how the star and detour graphs represent extreme values of the fixation probability. Here *s* = 0.005.

The star graphs and the detour graphs constitute limiting structures for the range of probabilities of fixation for undirected graphs under sufficiently weak selection (**Figure 5B**). The difference in probability of fixation between the detour graph and a well-mixed population is close to zero when graph size is small, however it decreases sharply as population size increases. This explains why strong suppressors are prevalent in large populations, but rarely observed in small populations. Although we did not rigorously prove that detour graphs serve as the lower bound for the probability of fixation under the Birth-death update, this is empirically observed in graphs of small size (Möller et al., 2019). It is reasonable to assume the existence of large graphs that have stronger suppression, but this only reinforces our point that suppressors are more prevalent in larger populations. The result is particularly biologically interesting. Imagine populations with individuals fixed in space, such as species in an ecosystem or cells in biological tissues. These spatial populations can be reasonably approximated by random geometric or Waxman graphs. As shown in the previous section, these types of populations are likely to be suppressors under the Birth-death update. If the size of the population were to decrease (for example, environmental catastrophes or injury and aging of tissues), not only will the force of drift increase in the population, but also the suppressive capability of the population against the invasion of beneficial mutation will be compromised. This could lead to increased likelihood of beneficial mutations propagating in the population (and potential rescue the population from extinction), or increased rate of accumulation of deleterious variants.

### Application to mutation accumulation in hematopoietic stem cell populations

We apply the theory developed above to the study of rates of mutation accumulation in the hematopoietic stem cell (HSC) population of the bone marrow. Hematopoietic stem cells reside in specialized micro-environments, or niches, where distinct mesenchymal cells, the vasculature, and differentiated hematopoietic cells interact to regulate stem cell maintenance and differentiation (Morrison and Scadden, 2014; Baccin et al., 2020). These niches are fixed in location, have heterogeneous spatial structure, and stem cells are in constant competition for niche occupancy (Celso and Scadden, 2011; Glait-Santar et al., 2015).

New innovations in imaging techniques and our ability to process these images at scale offer unprecedented opportunities to study the spatial heterogeneity of stem cell niches. Just as demographic surveys can reveal the rates at which a contagious disease can spread through a spatially heterogenous population, microscopy datasets allow us to quantify the patterns of cellular spatial variation and study how these topologies shape evolutionary dynamics. We build the networks of stem cell niches and use the inferred spatial topologies to infer the accumulation rate of driver mutations, the main cause of cancer in cycling tissues (Klein and Simons, 2011).

We use recent datasets that provide the spatial location of hematopoietic stem and progenitor cells in four samples of mouse tibia (Coutu et al., 2018) and the spatial location of CXCL12-abundant reticular cells (which critically modulate hematopoiesis at various levels, including hematopoietic stem cell maintenance), each with two images of two anatomically distinct regions (the diaphysis and the metaphysis), in total 16 cellular populations in 8 bone marrow samples (Gomariz et al., 2018). The presence of these cells types is a proxy for the spatial location of the hematopoietic stem cell niche. Live-animal tracking of individual hematopoietic stem cells shows that MFG cells, a largely quiescent population with long-term self-renewal capability, can migrate and displace across an average distance of 8.69*µ*m in a 2.5 hour period (Christodoulou et al., 2020). HSCs have a median replication time of 1.7 weeks (Abkowitz et al., 2000). During homeostasis, the rate of replication should balance the rate of depletion. This leads to an estimated interaction range of 1028*µ*m.

We build networks where every HSC niche constitutes a node in the graph and stem cell movement between niches is represented by edges between the nodes. An edge is added between two nodes if the distance between the two nodes is less than a cut-off radius, similar to the generation of a random geometric graph. The samples vary in dimensions, number of cells, and segmentation techniques. We normalize the data by expressing the distance in units of the average distance between shortest pairs of cells (62.72 *µ*m for HSC). An example of the resulting networks is shown in **Figure 6**. For illustration purposes, for the network shown, the cutoff distance is set to 300*µ*m (4.78× the distance between shortest pair distance). We also construct networks with a probabilistic connection function (Waxman, 1988), and observe no qualitative difference.

**Figure 6:**
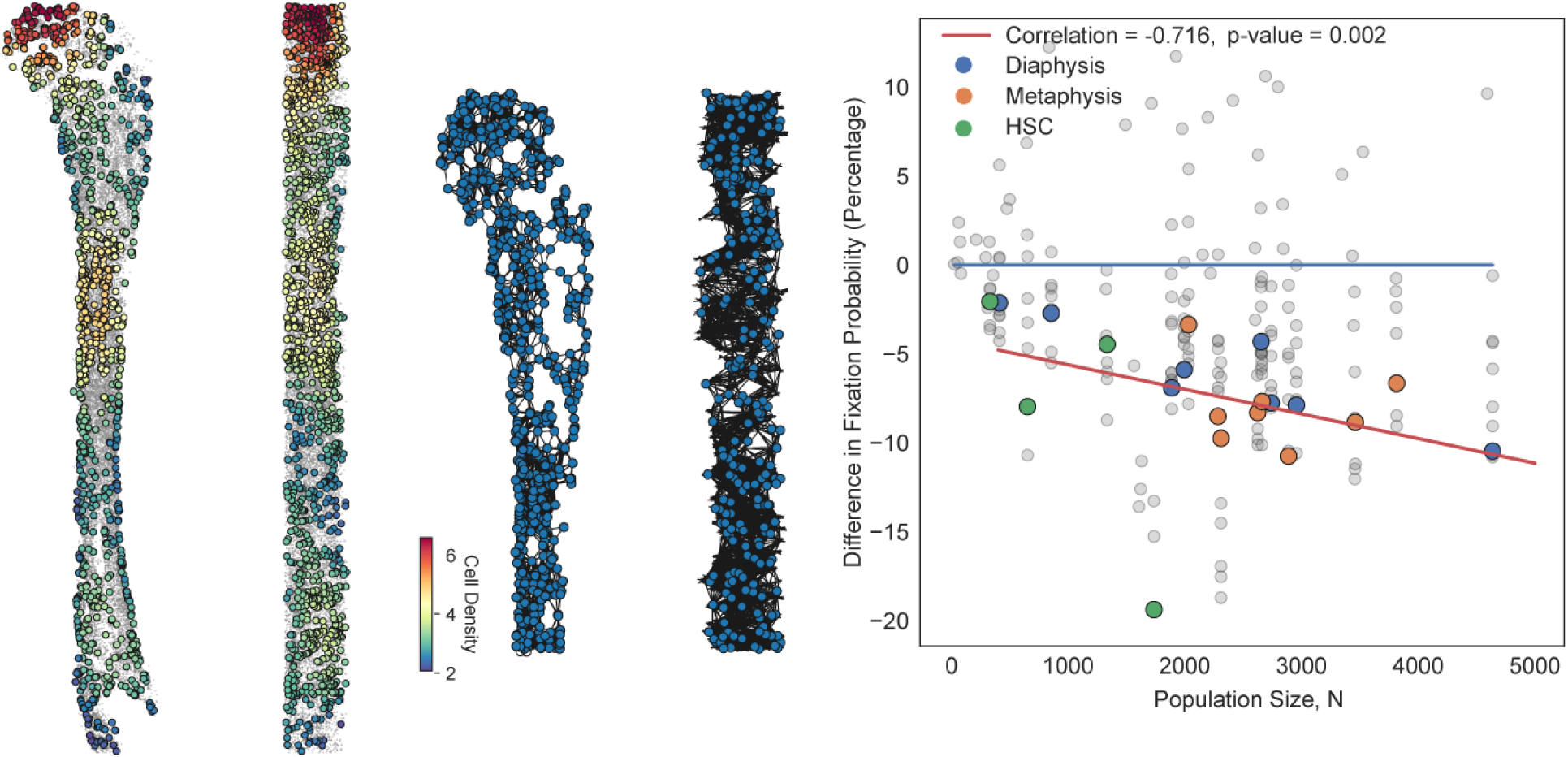
The spatial networks of the hematopoietic stem cell (HSC) populations. We use a public dataset of spatial locations in the bone marrow consisting of HSC locations (source data: (Coutu et al., 2018)) and proxy niche component locations (source data: (Gomariz et al., 2018)). On the left, color dots represent an example of the spatial locations of stem cells in the cell population, while the colors corresponds to the density of cells. We show an example of the spatial distribution of hematopoietic stem cells in the mouse tibia: the *xy* cross-section of the tibia with cells counted in a 260 *µ*m × 350 *µ*m area and the *yz* cross-section of the tibia with cells counted in a 350 *µ*m × 11.5 *µ*m area. Using this data, we build the resulting geometric random graphs (the cut-off radius of the illustration is set to 300 *µ*m for illustration purposes). On the right, we use these networks to study evolutionary dynamics on these stem cell niche spatial networks as a function of their population size. Here, the diaphysis is the shaft or central part of the tibia and the metaphysis is the neck portion of the bone (source data: (Gomariz et al., 2018)). The gray dots represent probabilities of fixation using cut-off distances ranging from 2 to 20 times the average distance between shortest pairs. The color dots use a cut-off distance of 15. Here, *s* = 0.005 and *Ns* varies with population size. We pick this selective advantage to model a mutation with moderately strong selection pressure.

We analyze evolutionary dynamics on networks generated using cut-off radii ranging from 2 to 20 times the average distance between shortest pairs. We interpret the cut-off distance as the maximum distance a potentially mutated HSC could travel in its entire lifespan. The estimated interaction range of 1028*µ*m corresponds to 16.4× the distance between the shortest pair distance. The difference in probabilities of fixation, compared to those in well-mixed populations, is plotted against the population size in **Figure 6**. The fixation probabilities are either close to or lower than that of the well-mixed. The networks with fixation probability above the well-mixed line are networks generated using extreme values of the edge cutoff, such as two times the shortest distance. The color dots are the results using a cut-off distance of 15, which is the closest to our estimated biological interaction range.

These graphs have a highly assortative mixing pattern, therefore, based on our results in previous sections, we find strong suppression of selection in the hematopoietic stem cell populations, invariant to the underlying update process. Our results also show that the strength of suppression increases as the stem cell population size increases. The fixation probability in populations constructed with the same cut-off radius shows a negative correlation with population size (Pearson correlation of −0.688 and p-value of 0.002). A similar conclusion is reached with most other cut-off distances (see examples of cut-off 10 and 20 in the **Supplementary Figure S3**). This observation could partially explain the observed reduction in cancer incidence in large organisms as stated by Peto’s paradox (Caulin and Maley, 2011). This also implies that processes such as injury or aging, that lead to reduced stem cell and niche count, could lead to the increased likelihood of beneficial mutations propagating in the population and an increased risk of developing cumulative diseases of aging, such as cancer.

## Discussion

Graphs represent a powerful tool to mathematically represent a population’s structure of spread or interaction and to ask how properties of this structure shape the balance of evolutionary forces. However, obtaining closed form solutions for evolutionary dynamics on graphs has been particularly difficult. Here we introduce new theoretical and computational methods to rigorously study the role of graph topology on shaping evolutionary dynamics. We focus on parameters of the degree distribution and the graph mixing pattern because these distributions inform on graph-wide properties of the fundamental building blocks of a network: the nodes and the edges.

Previous theoretical studies on the role of the degree distribution of the network of interaction in epidemiological studies assume that nodes of the same degree are fundamentally identical and fail to study properties of the distribution independent of each other (May and Lloyd, 2001; Pastor-Satorras et al., 2015; Sood and Redner, 2005; Antal et al., 2006). These models also tend to ignore the role of edge heterogeneity, by assuming that the networks are randomly mixed and the frequencies of edge types can be represented in terms of the frequencies of node degrees themselves. While this assumption leads to approximations that predict the relative probability of fixation of different initial mutant configurations, it nonetheless predicts that all graphs should have the same probability of fixation as the well-mixed population (Antal et al., 2006). Our work provides the first general theory for the role of network topology on evolutionary dynamics, analyzing both the contribution of properties of nodes and their distribution, as well as properties of the edges.

We show that the probability of fixation of a new mutation appearing on a random node can be approximated by solving a system of quadratic equations with number of variables depending on the number of degrees in the graph, which in most practical cases is efficient, even in large populations. By tuning the first moments of the degree distribution independently of each other, we analyze how the mean and variance in degree changes probabilities and times to fixation. For example, we show that the probability of fixation under the Birth-death update increases monotonically as a function of the variance of the degree distribution. This is because the parameter that controls degree heterogeneity also controls the mixing pattern of the graph, by changing the connection bias towards nodes of higher degree.

Moreover, we find the relevant selective parameter of suppression or amplification (*α*_*dB*_ and *α*_*Bd*_), predictive of whether the network is an amplifier or suppressor of the force of selection. While for the death-Birth process, this constant depends on properties of the degree distribution, for the Birth-death process, this constant is composed of parameters of both the mixing pattern and the degree distribution of the graph.

If degree distribution is held constant, increasing the amplification parameter corresponds to increasing the disassortativity of the graph. Or reversely, increasing the disassortativity of a network increases the probabilities of fixation monotonically across multiple random graph families. For the death-Birth process, increasing disassortativity also increases the fixation probability, but only when selection is larger than order of 1*/N*. For random graphs, it has been shown that the fixation probability correlates with the variance of vertex temperature, which is defined as the sum of the degrees inverse of all neighbors (Tan and Lü, 2014). Measures of temperature inherently encode both degree and edge information; thus, this is connected to the amplification parameter we derived. Our work goes beyond observed correlation and provides an analytical explanation of how the statistical properties of the nodes and edges shape fixation probabilities.

We also show that, contrary to prior empirical observations on small graphs (Hindersin and Traulsen, 2015), the Birth-death process can also be a strong suppressor of selection and not just an amplifier. For example, detour graphs are a class of graphs that show strong suppression under the Birth-death process. These graphs are extremely assortative and the magnitude of the suppression is shown to depend on the detour size. We analytically find the optimal suppression sizes of the detour graphs for any given population size, and empirically show that the magnitude of maximum suppression decreases with population size. Since the lower bound of fixation probability decreases as population size increases, there are hints at the possible existence of an arbitrarily strong suppressor that neglects selective advantages in a large population.

In biological settings, such as spatially complex ecosystems or cells in biological tissues, large populations are therefore more likely to be suppressors under the Birth-death update. If an event were to decrease the size of the population (for example, environmental catastrophes that lead to the destruction of forests or injury and aging of tissues), not only will the force of drift increase in the population, but also the suppressive capability of the population against the invasion of beneficial mutation will be compromised. One caveat is that this magnitude of suppression also depends on the strength of selection, and these topologies can transition from suppressor to amplifier given large enough selection pressure.

In rapidly cycling tissues, tissue maintenance and repair are coordinated by stem cells, which are routinely stochastically lost and replaced in the population (Klein and Simons, 2011) and thus instrumental for studying rates of evolution and mutation accumulation (Tomasetti et al., 2017). Models that approximate stem cell spatial structures as regular lattice networks have previously been used to explain results from lineage tracing experiments in the colon (Lopez-Garcia et al., 2010), epidermis (Mesa et al., 2018), and seminiferous tubules (Klein et al., 2010). Recent advances in quantitative imaging and single-cell sequencing technology allow for the high throughput acquisition of individual cells’ spatial location and genetic or cellular identity (Gomariz et al., 2018); Acar et al., 2015; Coutu et al., 2018; Codeluppi et al., 2018; Butler et al., 2018; Moffitt et al., 2018). The spatial relationship of individual cells is often represented as a network, and such representations have been instrumental in finding ligand-receptor pairs responsible for cell-cell communications (Yuan and Bar-Joseph, 2019), or identifying correlated expression of genes between neighboring cells (Dries et al., 2019).

By analyzing recent imaging data on the spatial organization of hematopoietic stem cells in the bone marrow, we find that stem cell populations are organized to minimize the chance of cancer initiation. Previous theoretical studies on population dynamics of marrow tissue either ignore the structure of the tissue, consider the topology of small populations of stem cells, or assume cells are arranged in lattices where every node has the same degree. Watson et al. (2020) applied population genetic models in well-mixed populations to estimate mutation accumulation in cells in blood from sequencing data and found that positive selection is a major force shaping clonal hematopoiesis. In the presence of clonal interference, lattice spatial structures also increase the waiting time for cancer, leading to a patchwork structure of non-uniformly sized clones (Martens et al., 2011). Hindersin et al. (2016b) et al proposed topologies for small populations (∼12 cells) of stem cells at the intestinal crypt that could suppress evolution in both Birth-death and death-Birth processes; however, the topology relies on weighted and directed edges. Here, we use recent imaging data to build the spatial structures of the stems cell niches in the bone marrow and find these structures to be strong suppressors of evolutionary dynamics (for both dB and Bd processes). Our work suggests when studying biological populations, samples should be drawn from regions where the connectivity is heterogeneous, since driver mutations occurring in these regions have a higher chance of spreading in the population.

Since networks are also the ideal mathematical proxy for population interaction and spread of cultural phenotypes, we collected Facebook friendship networks from recent datasets that span across 20 universities (Rossi and Ahmed, 2015) and observe that the evolutionary dynamics of culture is, in contrast to the stem cell networks, heavily dependent on the underlying process. Since most of the social networks exhibit community structure, we extract the connected communities from the network using a community structure detection algorithm (Newman, 2003). The data set we use therefore consists of 62 connected networks of sizes 150 − 1800 (**Supplementary Figure S4**). As expected, under the Birth-death process, these networks are mostly amplifiers, while suppressing selection under the death-Birth process. Although recent work suggests social networks weakly exhibit the scale-free property, where the degree distribution follows a power-law, these networks still tend to contain large heterogeneity in degree (Broido and Clauset, 2019). We do observe a weak suppressor (*n* = 1360) under the Birth-death update, and this suppression can be attributed to the assortative nature of the mixing pattern (*α*_*Bd*_ = 0.649, degree correlation equal to 0.122). These results highlight the importance of mixing patterns in inferring evolutionary dynamics on the network considered. A thorough theoretical understanding of the role of network topology on evolutionary dynamics makes many more questions ripe for exploration. For example, does the network topology affect patterns of mutational clonal interference? In other words, can the sweep on an initial mutation to fixation change the amplifier or suppressor property of a graph for a second mutation entering the population? Further work on how network properties shape evolutionary dynamics will also help us understand how to construct spatial structures in the limit of either suppression or amplification across various biological systems, natural or artificial. This would allow controlled suppression against the spread of unwanted variants and delay of population collapse. Reversely, we could also use population structure as a screening tool for faster amplification of newly discovered beneficial mutations or optimized protein complexes for medical or industrial applications.

## Acknowledgments

We gratefully acknowledge support from the United States-Israel Binational Science Foundation (award no. 2019266) and from the NIH T32 training grant (no. T32 EB009403). We also thank Daniel Coutu for help accessing part of the recent bone marrow imaging dataset used in the study. This research was done using resources provided by the Open Science Grid, which is supported by the National Science Foundation award 1148698, and the U.S. Department of Energy’s Office of Science.

