## Supplementary Material for "A theory of evolutionary dynamics on any complex spatial structure"

#### Notes on the graph rewiring algorithm

There are many graph generation algorithms across network families. Most of them allow for the tuning of network properties by changing a few input parameters. Since these algorithms are specific to a particular network family, one cannot continuously and smoothly explore the space of network properties and compare networks across different families. As described in the main text, we use simulated annealing procedures and degree preserving edge swapping to systematically tune networks properties across graph families. We use combinations of parameters in five graph families and study six main graph characteristics (mean degree, variance in degree, third moment of degree distribution , network modularity, average clustering, and network assortativity), predictive of evolutionary dynamics. We use PCA to reduce the dimensionality of the data (**Supplementary Figure S5**). We are able to capture 89% of the variation in network statistics using only the first three principle components. We observe that the graph families are clustered in this lower dimensional embedding space, with gaps in space existing between the network families. Therefore, our method allows us to sample graphs not accessible by traditional network generation algorithms specific to particular graph families. The black line in **Supplementary Figure S5** shows a continuous path through intermediate graphs, previously inaccessible using other methods, generated by the graph rewiring.

#### Analytic approximation for the death-Birth process

Here we present a full description of the analytic approach under the death-Birth update rule.

At every time step, let $T_n^+$ and $T_n^-$ denote the probabilities that node $n$ changes its allelic type towards or away from the mutant type. Let $x_n$ denote the frequency of the mutant at this node $n$ ($x_n = 1$ means a mutant occupies node $n$ and $x_n = 0$ means the node is occupied by the wild-type allele), $\mathcal{N}(n)$ denote the set of nodes connected to $n$ and degree $d_n$ denote the size of $\mathcal{N}(n)$. We can write

$$\begin{aligned} T_n^+ &= \frac{1-x_n}{N}(1+s)\frac{\sum_{m\in\mathcal{N}(n)} x_m}{d_n + s\sum_{m\in\mathcal{N}(n)} x_m} \\ T_n^- &= \frac{x_n}{N}\frac{\sum_{m\in\mathcal{N}(n)}(1-x_m)}{d_n + s\sum_{m\in\mathcal{N}(n)} x_m}. \end{aligned} \tag{1}$$

The $(1-x_n)/N$ term in $T_n^+$ corresponds to the probability that node $n$ is both a wild-type and is also selected to die. The rest of the terms in $T_n^+$ correspond to the probability that a neighboring mutant node is selected to replace node $n$ and can be written as the fraction of the mutant neighbor fitness over the total fitness of neighbors of node $n$. This makes $T_n^+$ and $T_n^-$ difficult to work with. Using a power series expansion we can write

$$\frac{1}{d_n + s\sum_{m\in\mathcal{N}(n)} x_m} = \frac{1}{d_n} - \frac{s}{d_n^2}\sum_{m\in\mathcal{N}(n)} x_m + s^2\mathcal{O}(x^2). \tag{2}$$

This will later make the calculations easier.

The approach we take here is to use the node degree distribution, and only keep track of the mutant frequencies $x_i$ at all $N_i$ nodes of the same degree $d_i$. Let $D = \{d_1, d_2, ..., d_i, ...\}$ represent the set of all possible node degrees. We denote the frequency of nodes of degree $d_i$ in the population by $p_i$. To model node degree mixing, we use $p_{ij}$ to denote the probability that a node of degree $d_i$ is connected to a node of degree $d_j$. The probability that the mutant frequency increases by $1/N_i$ in nodes of degree $d_i$, $T_i^+$, is given by

$$
\begin{aligned}
T_i^+ &= (1+s)\sum_{n\in G}\left[\delta(d_i,d_n)\frac{1-x_i}{N}\left(\sum_{m\in\mathcal{N}(n)} x_m\right)\left(\frac{1}{d_i}-\frac{s}{d_i^2}\sum_{m\in\mathcal{N}(n)} x_m + s^2\mathcal{O}(x^2)\right)\right]\\
&= (1+s)\frac{1-x_i}{N}\sum_{n\in G}\left[\delta(d_i,d_n)\left(\sum_{j\in D} e_{nj}x_j\right)\left(\frac{1}{d_i}-\frac{s}{d_i^2}\sum_{j\in D} e_{nj}x_j + s^2\mathcal{O}(x^2)\right)\right]\\
&= (1+s)\left[\frac{1-x_i}{N}\frac{1}{d_i}\sum_{n\in G}\delta(d_i,d_n)\sum_{j\in D} e_{nj}x_j + \frac{1}{N}\frac{s}{d_i^2}\sum_{n\in G}\delta(d_i,d_n)\left(\sum_{j\in D} e_{nj}x_j\right)^2\right] + s^2\mathcal{O}(x^3)\\
&= (1+s)\frac{1-x_i}{N}\frac{1}{d_i}\sum_{j\in D} e_{ij}x_j + s\mathcal{O}(x^2) + s^2\mathcal{O}(x^3)\\
&= (1+s)\frac{1-x_i}{N}\frac{1}{d_i}\sum_{j\in D} Np_ip_{ij}d_ix_j + s\mathcal{O}(x^2) + s^2\mathcal{O}(x^3)\\
&= (1+s)(1-x_i)\sum_{j\in D} p_ip_{ij}x_j + s\mathcal{O}(x^2) + s^2\mathcal{O}(x^3), \qquad (3)
\end{aligned}
$$

while the probability that the mutant frequency decreases by $1/N_i$, $T_i^-$, is given by

$$
T_i^- = x_i\sum_{j\in D} p_ip_{ij}(1-x_j) + s\mathcal{O}(x^2) + s^2\mathcal{O}(x^3). \qquad (4)
$$

Here, $\delta(d_i,d_n)$ is the Kronecker delta function, the set $G$ represents all the nodes in the graph, $e_{nj}$ denotes the number of edges that connect node $n$ to nodes of degree $d_j$ and $e_{ij}$ denotes the number of edges that connect nodes of degree $d_i$ to nodes of degree $d_j$.

The probability of fixation of allele $a$ can then be approximated using the diffusion approximation. We will first need to calculate the first and second moment of the change in frequency of the mutant allele at all nodes of degree $d_i$, at every time step:

$$
\begin{aligned}
E[\Delta x_i] &= (T_i^+ - T_i^-)\Delta x_i = \frac{1}{Np_i}(T_i^+ - T_i^-) = \mathcal{O}(x), &(5)\\
E[(\Delta x_i)^2] &= (T_i^+ + T_i^-)(\Delta x_i)^2 = \frac{1}{N^2p_i^2}(T_i^+ + T_i^-) = \mathcal{O}(x), &(6)\\
E[\Delta x_i\Delta x_j] &= 0, &(7)\\
E[\Delta x_i]E[\Delta x_j] &= (T_i^+ - T_i^-)(T_j^+ - T_j^-)(\Delta x_i)^2 = \frac{1}{N^2p_ip_j}(T_i^+ - T_i^-)(T_j^+ - T_j^-) = \mathcal{O}(x^2). &(8)
\end{aligned}
$$

This allows us to write the mean change in mutant frequency at every time step as

$$\mu_i = \frac{E[\Delta x_i]}{\Delta t} = \frac{1}{p_i}(T_i^+ - T_i^-). \tag{9}$$

It is worth noting that in many diffusion models the variance can be approximated as the second moment and the covariance is omitted since the product of the first moments is often on the order of $s^2$. This is not the case in our model, since here the first moment is on the order of $s^0$ therefore the product of the first moments does not go away by assuming sufficiently small $s$.

The variance in mutant frequency change can be written as

$$\sigma_{ii} = \frac{E[(\Delta x_i)^2] - (E[\Delta x_i])^2}{\Delta t} = \frac{1}{Np_i^2}[T_i^+ - T_i^- - (T_i^+ - T_i^-)^2], \tag{10}$$

while the covariance can be written as

$$\sigma_{ij} = \frac{-E[\Delta x_i]E[\Delta x_j]}{\Delta t} = -\frac{1}{Np_ip_j}(T_i^+ - T_i^-)(T_j^+ - T_j^-). \tag{11}$$

We write the Kolmogorov backward equation

$$\begin{aligned}\frac{\partial P}{\partial t} &= \frac{1}{2}\sum_{i,j\in D}\sigma_{ij}\frac{\partial^2 P}{\partial x_i \partial x_j} + \sum_{i\in D}\mu_i \frac{\partial P}{\partial x_i} \\ &= -\frac{1}{2N}\sum_{i\neq j}\mathcal{O}(x^2)\frac{\partial^2 P}{\partial x_i \partial x_j} + \sum_{i\in D}\mathcal{O}(x)\left(\frac{1}{2N}\frac{\partial^2 P}{\partial x_i^2} + \frac{\partial P}{\partial x_i}\right),\end{aligned} \tag{12}$$

and solve for zero

$$-\frac{1}{2N}\sum_{i\neq j}\mathcal{O}(x^2)\frac{\partial^2 P}{\partial x_i \partial x_j} + \sum_{i\in D}\mathcal{O}(x)\left(\frac{1}{2N}\frac{\partial^2 P}{\partial x_i^2} + \frac{\partial P}{\partial x_i}\right) = 0. \tag{13}$$

Given the initial mutant frequencies $\vec{x}$, $P(\vec{x})$ gives an approximation for the fixation probability of the mutant allele $a$. It is difficult to find a closed form solution for $P(\vec{x})$, since coefficients in the PDE in equation (13) are polynomials of $x$. Due to the similarity between the Kolmogorov backward equation here and the Kolmogorov backward equation for the finite island model (Tachida and Iizuka, 1991), we can use singular perturbation methods to approximate the solution (Gavrilets and Gibson, 2002). This method tries to find the solution to the PDE of interest near singular points, where the function changes value rapidly. This usually occurs in the region of space where the PDE coefficients vanish and therefore where the first derivatives are large in magnitude.

For our PDE, the singular points occur at $\vec{x} = \vec{0}$ and $\vec{x} = \vec{1}$. For $s > 0$, we solve the PDE at $\vec{x} = \vec{0}$, while for $s \leq 0$, we solve for $\vec{x} = \vec{1}$. Intuitively, the fixation probability for any mutant with selective advantage $s$ should be unity in the deterministic infinite population case.

In finite populations however, fixation is controlled by both the force of selection and the force of drift. The force of drift is proportional to $1/N$ and can cause even beneficial mutants to become extinct. As mutant frequency increases in the population, past establishment, the force of selection starts to dominate the force of drift and the fixation probability starts approaching one rapidly. For deleterious mutations, the fixation probability should be small unless the number of mutants is close to population size $N$; therefore, for $s \leq 0$, $P$ decreases to 0 when $\vec{x}$ moves away from $\vec{1}$.

For $s > 0$, we introduce new variables $y_i$, such that $\epsilon y_i = x_i$, where $\epsilon = \frac{1}{N}$. We can write

$$\frac{\partial P}{\partial x_i} = \frac{\partial P}{\partial y_i}\frac{dy_i}{dx_i} = \frac{1}{\epsilon}\frac{\partial P}{\partial y_i} \tag{14}$$

and

$$\frac{\partial^2 P}{\partial x_i \partial x_j} = \frac{\partial}{\partial x_i}\left(\frac{\partial P}{\partial y_j}\frac{dy_j}{dx_j}\right) = \frac{\partial^2 P}{\partial y_i \partial y_j}\frac{dy_i}{dx_i}\frac{dy_j}{dx_j} + \frac{\partial P}{\partial y_j}\frac{\partial^2 y_i}{\partial x_i \partial x_j} = \frac{1}{\epsilon^2}\frac{\partial^2 P}{\partial y_i \partial y_j}. \tag{15}$$

We can substitute (14) and (15) into (13) and write

$$-\frac{1}{2}\sum_{i \neq j}\epsilon^{-1}\mathcal{O}(\epsilon^2 y^2)\frac{\partial^2 P}{\partial y_i \partial y_j} + \sum_{i \in D}\mathcal{O}(\epsilon y)\left(\frac{1}{2}\epsilon^{-1}\frac{\partial^2 P}{\partial y_i^2} + \epsilon^{-1}\frac{\partial P}{\partial y_i}\right) = 0. \tag{16}$$

For large population sizes, $\epsilon = 1/N$ becomes vanishingly small, therefore in the equation above, we can ignore higher order terms of $\epsilon$. Therefore we can approximate (16) by

$$\sum_{i \in D}\mathcal{O}(y)\left(\frac{1}{2}\frac{\partial^2 P}{\partial y_i^2} + \frac{\partial P}{\partial y_i}\right) = 0. \tag{17}$$

We exand equation (17) and write

$$\sum_{i,j \in D} p_i p_{ij}\left(\frac{1}{2p_i^2}((1+s)y_j + y_i)\frac{\partial^2}{\partial y_i^2} + \frac{1}{p_i}((1+s)y_j - y_i)\frac{\partial}{\partial y_i}\right)P = 0 \tag{18}$$

It is important to note that the Kolmogorov backward equations for the death-Birth model we consider here and the Death-birth voter model (the update process where a node is first picked for death with probability inversely proportional to fitness, and a random neighbor is then selected to replace it) are identical after singular perturbation. The Kolmogorov backward equations for the Birth-death model considered here and the birth-Death model also share the same equations. This implies that the dB and Db should have identical fixation probabilities for the same network. Indeed, the two processes lead to similar evolutionary dynamics (Chen et al., 2013).

The solution to the differential equation in (18) has the form

$$P = c_0 + c_1 \exp\Big\{ -\sum_j p_j A_j y_j \Big\}. \tag{19}$$

We can substitute this solution back into the Kolmogorov backward equation (18) and solve for the unknown exponents:

$$\begin{aligned}&\sum_{i,j\in D}\left(\frac{1}{2}(1+s)A_i^2 p_i p_{ij} y_j + \frac{1}{2}A_i^2 p_i p_{ij} y_i - (1+s)A_i p_i p_{ij} y_j + A_i p_i p_{ij} y_i\right)\\ &= \sum_{i,j\in D}\left(\frac{1}{2}(1+s)A_j^2 p_j p_{ji} y_i + \frac{1}{2}A_i^2 p_i p_{ij} y_i - (1+s)A_j p_j p_{ji} y_i + A_i p_i p_{ij} y_i\right) = 0.\end{aligned} \tag{20}$$

We end up with the following system of quadratic equations to solve:

$$\sum_{j\in D}\Big((1+s)A_j^2 p_j p_{ji} + A_i^2 p_i p_{ij} - 2(1+s)A_j p_j p_{ji} + 2A_i p_i p_{ij}\Big) = 0 \quad \forall i. \tag{21}$$

This is a system of $|D|$ (the number of unique degrees in the graph) elliptic equations in $|D|$-dimensional space and the solution to this system corresponds to the set of points in space where all these surfaces intersect. There is a trivial intersection point at the origin. This solution, however, causes $P$ to be undefined so it is not the solution we are interested in. Assuming there is a non-trivial real solution to this system, we can use geometric intuition to estimate where the solution is. We do this by summing all the equations in the system to get the following equation

$$\begin{aligned}&\sum_{i,j\in D}\Big((1+s)A_j^2 p_j p_{ji} + A_i^2 p_i p_{ij} - 2(1+s)A_j p_j p_{ji} + 2A_i p_i p_{ij}\Big)\\ &= \sum_{i,j\in D}\Big((1+s)A_i^2 p_i p_{ij} + A_i^2 p_i p_{ij} - 2(1+s)A_i p_i p_{ij} + 2A_i p_i p_{ij}\Big)\\ &= \sum_{i\in D}\Big((1+s)A_i^2 p_i + A_i^2 p_i - 2(1+s)A_i p_i + 2A_i p_i\Big)\\ &= \sum_{i\in D}\Big[(2+s)A_i^2 p_i - 2sA_i p_i\Big] = 0.\end{aligned} \tag{22}$$

In elliptic form,

$$\sum_{i\in D} p_i \left(A_i - \frac{s}{2+s}\right)^2 = \left(\frac{s}{2+s}\right)^2. \tag{23}$$

This equation provides valuable information on the dynamics of the system. This ellipsoid contains all solutions to the system since it is constructed from a linear combination of these ellipsoids. It is centered at $s/(2+s)\vec{1}$, with axial lengths proportional to $s/(2+s)$. In the neutral case where $s = 0$, this ellipsoid collapses into a point at the origin. Since all solutions of the elliptic system coincide with this point, the system has exactly one real solution at the origin. When $s$ increases from 0, the distance between the solution at the origin and all other real solutions grows proportional to the axial lengths, which themselves are proportional to $s/(2+s)$. We will use these intuitions later to derive simpler forms of the solutions of the entire system.

Next, we use regular perturbation to study the system. We can write the solutions of the system as

$$A_i = A_{i,0} + sA_{i,1} + \mathcal{O}(s^2) \tag{24}$$

Substitute this and the following

$$A_i^2 = A_{i,0}^2 + sA_{i,0}A_{i,1} + \mathcal{O}(s^2) \tag{25}$$

into the elliptic system and we have

$$\sum_{j\in D} \Big[(1+s)(A_{j,0}^2 + sA_{j,0}A_{j,1} - 2A_{j,0} - 2sA_{j,1})p_j p_{ji} + (A_{i,0}^2 + sA_{i,0}A_{i,1} + 2A_{i,0} + 2sA_{i,1})p_i p_{ij}\Big] = \mathcal{O}(s^2). \tag{26}$$

In the order of $s^0$, we can derive $A_{i,0}$ using

$$\sum_{j\in D} \Big[(A_{j,0}^2 - 2A_{j,0})p_j p_{ji} + (A_{i,0}^2 + 2A_{i,0})p_i p_{ij}\Big] = 0. \tag{27}$$

This is exactly the elliptic system corresponding to the neutral case where $s = 0$. We know from the argument above that this system only has one real solution at the origin.

For the order of $s^1$, we can derive $A_{i,1}$ using

$$\sum_{j\in D} \Big( -2A_{j,1}p_j p_{ji} + 2A_{i,1}p_i p_{ij}\Big) = 0. \tag{28}$$

Using the fact that the number of edges going from nodes of degree $d_i$ to nodes of degree $d_j$ is equal to the number of edges going from nodes of degree $d_j$ to nodes of degree $d_i$ (the handshaking lemma), we can write

$$p_i p_{ij} d_i = p_j p_{ji} d_j. \tag{29}$$

We rewrite the above as

$$\sum_{j\in D} p_j p_{ji}\Big( - A_{j,1} + A_{i,1}\frac{d_j}{d_i}\Big) = 0. \tag{30}$$

It follows that points on the line $A_{i,1} = Ad_i$ satisfy this equation. Substituting in (24), we now have an approximation of the solution of the elliptic system

$$A_i = sAd_i + \mathcal{O}(s^2). \tag{31}$$

This agrees with the fact that the real solutions of the elliptic system grow proportional to $s/(2+s)$ (from (23)).

We still have to find $A$. Since we know the solution to the system must also satisfy equation (23), the value of $A$ that approximates the solution of the system is

$$\begin{aligned}
&\sum_{i\in D}\Big[(2+s)p_i A^2 d_i^2 - 2sp_i A d_i\Big] = 0\\
&\implies (2+s)A^2\sum_{i\in D} p_i d_i^2 - 2sA\sum_{i\in D} p_i d_i = 0\\
&\implies (2+s)A^2\langle d^2\rangle - 2sA\langle d\rangle = 0\\
&\implies (2+s)A\langle d^2\rangle = 2s\langle d\rangle\\
&\implies A = \frac{2s}{2+s}\frac{\langle d\rangle}{\langle d^2\rangle}.
\end{aligned} \tag{32}$$

Here $\langle d^k\rangle$ represents the $k$-th moment of the degree distribution. We substitute this back into (19) and can thus find the constants that satisfy the boundary conditions that $P(\vec{x} = \vec{0}) = 0$ and $P(\vec{x} = \vec{1}) = 1$.

We can therefore write the approximation for the fixation probability as

$$\begin{aligned}P(\vec{x}) &= \left[1-\exp\left\{-N\frac{2s}{2+s}\frac{\langle d\rangle}{\langle d^2\rangle}\sum_{i\in D}p_i d_i x_i\right\}\right]\left[1-\exp\left\{-N\frac{2s}{2+s}\frac{\langle d\rangle}{\langle d^2\rangle}\sum_{i\in D}p_i d_i\right\}\right]^{-1}\\ &= \left[1-\exp\left\{-N\frac{2s}{2+s}\frac{\langle d\rangle}{\langle d^2\rangle}\sum_{i\in D}p_i d_i x_i\right\}\right]\left[1-\exp\left\{-N\frac{2s}{2+s}\frac{\langle d\rangle^2}{\langle d^2\rangle}\right\}\right]^{-1}. \qquad (33)\end{aligned}$$

Assuming the probability that the mutant was introduced uniformly into the network, the fixation probability is

$$P\left(\vec{x}=\frac{\vec{1}}{N}\right) = \left[1-\exp\left\{-\frac{2s}{2+s}\frac{\langle d\rangle^2}{\langle d^2\rangle}\right\}\right]\left[1-\exp\left\{-N\frac{2s}{2+s}\frac{\langle d\rangle^2}{\langle d^2\rangle}\right\}\right]^{-1}. \qquad (34)$$

To summarize, the fixation probability for the death-Birth process on a network is given by

$$P_{dB} = \frac{1-e^{-\alpha_{dB}s/(1+s/2)}}{1-e^{-\alpha_{dB}Ns/(1+s/2)}} \quad \text{where } \alpha_{dB} = \frac{\langle d\rangle^2}{\langle d^2\rangle} \qquad (35)$$

For the special case of uncorrelated networks, our approximation coincides with the fixation probability of the Death-birth voter model (Antal et al., 2006). As mentioned before, this is expected, as the Kolmogorov backward equations after singular perturbation are identical for the Death-birth and the death-Birth update rules. Our result however apply across network families, not just for the special case of uncorrelated networks.

In **Supplementary Figure S6** we show how well (35) approximates the fixation probability obtained from solving (21) numerically. In our derivation of the approximation, we ignored the $\mathcal{O}(s^2)$ portion of the roots of (21). The error that accumulates is on the order of $Ns^2$, therefore as long as $s << N^{-1/2}$ the approximation should hold. The approximate solution to the KBE remains accurate with few exceptions. In evolving populations, we are often interested in cases where there exists an interplay between drift and selection. This requires both forces to have similar magnitudes. This implies $s \approx \frac{1}{N}$, which implies $Ns^2 \approx \frac{1}{N} << 1$ in large populations.

#### Analytic approximation for the Birth-death process

We now discuss the probability of fixation of a new mutant under the Birth-death process. Following similar steps as the previous section, we start by writing down the probabilities $T_n^+$ and $T_n^-$ that a node $n$ switches allelic type towards or away from the mutant state. We can write

$$\begin{aligned} T_n^+ &= (1+s)\frac{\sum_{m\in\mathcal{N}(n)} x_m d_m^{-1}(1-x_n)}{N+s\sum_{m\in\mathcal{N}(n)} x_m} \\ T_n^- &= \frac{\sum_{m\in\mathcal{N}(n)}(1-x_m)d_m^{-1}x_n}{N+s\sum_{m\in\mathcal{N}(n)} x_m}. \end{aligned} \tag{36}$$

The denominator in $T_n^+$ is the total fitness of the population. Since it is shared across all $T$s we will represent it as $Nw$, where $w$ is the mean fitness of the population. The $x_m$ term in $T_n^+$ divided by the denominator corresponds to the probability that the focal node $n$ has a mutant neighbor node selected to reproduce for the Birth step. The rest of the terms in $T_n^+$ constitute the probability that node $n$ is the node selected at the death step. This probability of death is one over the degree of node $m$, an arbitrary neighbor of $n$. It might seem that this is as complicated as the transition probabilities for the dB update rule, and we should simplify using the power series. However, we do not need to do that here since the denominator can be multiplied out.

Similarly to the case of the death-Birth process, we use the degree mean field approximation. The probability that the mutant frequency increases by $1/N_i$ for nodes of degree $d_i$, $T_i^+$, is given by

$$\begin{aligned} T_i^+ &= \frac{(1+s)}{Nw}\sum_{n\in G}\left[\delta(d_i,d_n)\left(\sum_{m\in\mathcal{N}(n)} x_m d_m^{-1}(1-x_i)\right)\right] \\ &= \frac{(1+s)}{Nw}\sum_{n\in G}\left[\delta(d_i,d_n)\left(\sum_{j\in D} e_{jn}x_j d_j^{-1}(1-x_i)\right)\right] \\ &= \frac{(1+s)}{Nw}\sum_{j\in D} e_{ji}x_j d_j^{-1}(1-x_i) \\ &= \frac{(1+s)}{Nw}\sum_{j\in D} Np_jp_{ji}d_jx_jd_j^{-1}(1-x_i) \\ &= \frac{(1+s)}{w}\sum_{j\in D} p_jp_{ji}x_j(1-x_i), \end{aligned} \tag{37}$$

and the probability that the mutant frequency decreases by $1/N_i$ for nodes of degree $d_i$ , $T_i^-$ is

$$T_i^- = \frac{(1+s)}{w} \sum_{j \in D} p_j p_{ji} (1 - x_j) x_i. \tag{38}$$

Here, $e_{jn}$ denotes the number of edges that connect nodes of degree $d_j$ to $n$ and $e_{ji}$ denotes the number of edges that connect nodes of degree $d_j$ to nodes of degree $d_i$.

To write out the diffusion equation, we first need to compute the first and second moment of the change in frequency of the mutant allele at all nodes of degree $d_i$, at every time step:

$$E[\Delta x_i] = (T_i^+ - T_i^-)\Delta x_i = \frac{1}{Np_i}(T_i^+ - T_i^-) = w^{-1}\mathcal{O}(x) \tag{39}$$

$$E[(\Delta x_i)^2] = (T_i^+ + T_i^-)(\Delta x_i)^2 = \frac{1}{N^2 p_i^2}(T_i^+ + T_i^-) = w^{-1}\mathcal{O}(x) \tag{40}$$

$$E[\Delta x_i \Delta x_j] = 0 \tag{41}$$

$$E[\Delta x_i]E[\Delta x_j] = (T_i^+ - T_i^-)(T_j^+ - T_j^-)(\Delta x_i)^2 = \frac{1}{N^2 p_i p_j}(T_i^+ - T_i^-)(T_j^+ - T_j^-) = w^{-1}\mathcal{O}(x^2). \tag{42}$$

The mean change in mutant frequency at every time step can then be written as

$$\mu_i = \frac{E[\Delta x_i]}{\Delta t} = \frac{1}{p_i}(T_i^+ - T_i^-). \tag{43}$$

The variance can be written as

$$\sigma_{ii} = \frac{E[(\Delta x_i)^2] - (E[\Delta x_i])^2}{\Delta t} = \frac{1}{Np_i^2}[T_i^+ - T_i^- - (T_i^+ - T_i^-)^2]. \tag{44}$$

The covariance can be written as

$$\sigma_{ij} = \frac{-E[\Delta x_i]E[\Delta x_j]}{\Delta t} = -\frac{1}{Np_i p_j}(T_i^+ - T_i^-)(T_j^+ - T_j^-). \tag{45}$$

We can now write the Kolmogorov backward equation. Instead of substituting and writing all the coefficients in the equation, we are going to denote the the terms by their lowest degree of $x$

$$\begin{aligned}\frac{\partial P}{\partial t} &= \frac{1}{2}\sum_{i,j \in D} \sigma_{ij} \frac{\partial^2 P}{\partial x_i \partial x_j} + \sum_{i \in D} \mu_i \frac{\partial P}{\partial x_i} \\ &= -\frac{1}{2N}\sum_{i \neq j} w^{-1}\mathcal{O}(x^2)\frac{\partial^2 P}{\partial x_i \partial x_j} + \sum_{i \in D} w^{-1}\mathcal{O}(x)\left(\frac{1}{2N}\frac{\partial^2 P}{\partial x_i^2} + \frac{\partial P}{\partial x_i}\right).\end{aligned} \tag{46}$$

We are interested in the stationary solution where

$$-\frac{1}{2N}\sum_{i\neq j}\mathcal{O}(x^2)\frac{\partial^2 P}{\partial x_i \partial x_j}+\sum_{i\in D}\mathcal{O}(x)\left(\frac{1}{2N}\frac{\partial^2 P}{\partial x_i^2}+\frac{\partial P}{\partial x_i}\right)=0. \tag{47}$$

Note that we multiplied by the mean fitness $w$ on both sides to remove it from the PDE. By solving for $P(\vec{x})$, we have an approximation for the fixation probability given the initial mutant frequencies $\vec{x}$. Similarly as above, we apply singular perturbation to solve this system.

For $s>0$, we introduce new variables $y_i$, such that $\epsilon y_i = x_i$, where $\epsilon = \frac{1}{N}$. Substitute (14) and (15) into (47) and write

$$-\frac{1}{2}\sum_{i\neq j}\epsilon^{-1}\mathcal{O}(\epsilon^2 y^2)\frac{\partial^2 P}{\partial y_i \partial y_j}+\sum_{i\in D}\mathcal{O}(\epsilon y)\left(\frac{1}{2}\epsilon^{-1}\frac{\partial^2 P}{\partial x_i^2}+\epsilon^{-1}\frac{\partial P}{\partial x_i}\right)=0. \tag{48}$$

Ignoring vanishingly small higher-order terms of $\epsilon$, we write out terms of order $\epsilon^0$

$$\sum_{i\in D}\mathcal{O}(y)\left(\frac{1}{2}\frac{\partial^2 P}{\partial y_i^2}+\frac{\partial P}{\partial y_i}\right)=0. \tag{49}$$

Equation (49) can be expanded and written as

$$\sum_{i,j\in D}p_j p_{ji}\left(\frac{1}{2p_i^2}((1+s)y_j+y_i)\frac{\partial^2}{\partial y_i^2}+\frac{1}{p_i}((1+s)y_j-y_i)\frac{\partial}{\partial y_i}\right)P=0. \tag{50}$$

The solution to this differential equation has the form

$$P=c_0+c_1\exp\Big\{-\sum_j p_j A_j y_j\Big\}. \tag{51}$$

We can substitute this solution into the Kolmogorov backward equation (50) and solve for the unknown exponents:

$$\begin{aligned}&\sum_{i,j\in D}\left(\frac{1}{2}(1+s)A_i^2 p_j p_{ji} y_j+\frac{1}{2}A_i^2 p_j p_{ji} y_i-(1+s)A_i p_j p_{ji} y_j + A_i p_j p_{ji} y_i\right)\\ &=\sum_{i,j\in D}\left(\frac{1}{2}(1+s)A_j^2 p_i p_{ij} y_i+\frac{1}{2}A_i^2 p_j p_{ji} y_i-(1+s)A_j p_i p_{ij} y_i + A_i p_j p_{ji} y_i\right)=0. \end{aligned}\tag{52}$$

We end up with the following system of quadratic equations to solve

$$\sum_{j\in D}\Big((1+s)A_j^2p_ip_{ij}+A_i^2p_jp_{ji}-2(1+s)A_jp_ip_{ij}+2A_ip_jp_{ji}\Big)=0\quad\forall i. \tag{53}$$

Assuming there is a non-trivial real solution to this system, similarly as above, for the death-birth process, we can use geometric intuition to estimate where the solution is. We do so by summing all the equations in system to get the following equation

$$\begin{aligned}&\sum_{i,j\in D}\Big((1+s)A_j^2p_ip_{ij}+A_i^2p_jp_{ji}-2(1+s)A_jp_ip_{ij}+2A_ip_jp_{ji}\Big)\\&=\sum_{i,j\in D}\Big((1+s)A_i^2p_jp_{ji}+A_i^2p_jp_{ji}-2(1+s)A_ip_jp_{ji}+2A_ip_jp_{ji}\Big)\\&=\sum_{i\in D}\left[((1+s)A_i^2+A_i^2-2(1+s)A_i+2A_i)\sum_{j\in D}p_jp_{ji}\right]\\&=\sum_{i\in D}\left([(2+s)A_i^2-2sA_i]\sum_{j\in D}p_jp_{ji}\right).\end{aligned} \tag{54}$$

In elliptic form, we can write

$$\sum_{i\in D}\left[\left(A_i-\frac{s}{2+s}\right)^2\sum_{j\in D}p_jp_{ji}\right]=\left(\frac{s}{2+s}\right)^2\sum_{i\in D}\sum_{j\in D}p_jp_{ji}. \tag{55}$$

Like in the case of the death-Birth process, this equation provides valuable information on the dynamics of the system of ellipsoids. This ellipsoid contains all solutions to the system, since it is constructed from linear combinations of these ellipsoids. It is centered at $s/(2+s)\vec{1}$ with axial lengths proportional to $s/(2+s)$. In the neutral case where $s=0$, this ellipsoid collapses into a single point at the origin. Since all solutions of the elliptic system satisfy the equations, the system has exactly one real solution at the origin. As the strength of selection $s$ increases, the distance between the solution at the origin and all other real solutions grows proportional to the axial lengths, which themselves are proportional to $s/(2+s)$.

Next, we use regular perturbation to study the elliptic system. We can write the solution of the system as

$$A_i=A_{i,0}+sA_{i,1}+\mathcal{O}(s^2). \tag{56}$$

Substitute (56) and the following

$$A_i^2 = A_{i,0}^2 + sA_{i,0}A_{i,1} + \mathcal{O}(s^2) \tag{57}$$

into the system and we can write

$$\sum_{j\in D}\Big[(1+s)(A_{j,0}^2 + sA_{j,0}A_{j,1} - 2A_{j,0} - 2sA_{j,1})p_j p_{ji} \\ + (A_{i,0}^2 + sA_{i,0}A_{i,1} + 2A_{i,0} + 2sA_{i,1})p_i p_{ij}\Big] = \mathcal{O}(s^2). \tag{58}$$

In the order of $s^0$, we can derive $A_{i,0}$ using

$$\sum_{j\in D}\Big[(A_{j,0}^2 - 2A_{j,0})p_i p_{ij} + (A_{i,0}^2 + 2A_{i,0})p_j p_{ji}\Big] = 0. \tag{59}$$

This is exactly the elliptic system corresponding to the neutral case where $s = 0$. We know that this system only has one real solution at the origin.

For the order of $s^1$, we can derive $A_{i,1}$ using

$$\sum_{j\in D}\Big( - 2A_{j,1}p_i p_{ij} + 2A_{i,1}p_j p_{ji}\Big) = 0. \tag{60}$$

Using the handshaking lemma, (see (29)), we can rewrite the above as

$$\sum_{j\in D} p_j p_{ji}\Big( - A_{j,1}\frac{d_j}{d_i} + A_{i,1}\Big) = 0. \tag{61}$$

It follows that points on the line $A_i = Ad_i$ satisfy this equation. We now have an approximation of the solution of the system

$$A_i = sAd_i^{-1} + \mathcal{O}(s^2). \tag{62}$$

This agrees with the fact that real solutions grow proportional to $s/(2+s)$. Since we know the solution to the system must satisfy (55), we can find the intersection of (62) with (55) and the error from this point to the real intersection is of $\mathcal{O}(s^2)$.

The value $A$ that approximates the solution of the system is

$$\begin{aligned}
&\sum_{i\in D}\left([(2+s)A^2 d_i^{-2} - 2sAd_i^{-1}]\sum_{j\in D} p_j p_{ji}\right) = 0 \\
&\implies (2+s)A^2 \sum_{i,j\in D} p_j p_{ji} d_i^{-2} - 2sA \sum_{i,j\in D} p_j p_{ji} d_i^{-1} = 0 \\
&\implies (2+s)A \sum_{i,j\in D} p_j p_{ji} d_i^{-2} = 2s \sum_{i,j\in D} p_j p_{ji} d_i^{-1} \\
&\implies A = \frac{2s}{2+s}\left(\sum_{i,j\in D} p_j p_{ji} d_i^{-1}\right)\left(\sum_{i,j\in D} p_j p_{ji} d_i^{-2}\right)^{-1}. \qquad (63)
\end{aligned}$$

Substituting this back into (51) to find the constants that satisfy the boundary condition $p(\vec{x} = \vec{0}) = 0$ and $p(\vec{x} = \vec{1}) = 1$, we get the approximation for the fixation probability as

$$\begin{aligned}
P(\vec{x}) &= \frac{1 - \exp\Big\{ - NA \sum_{i\in D} p_i d_i^{-1} x_i \Big\}}{1 - \exp\Big\{ - NA \sum_{i\in D} p_i d_i^{-1} \Big\}} \\
&= \frac{1 - \exp\Big\{ - NA \sum_{i\in D} p_i d_i^{-1} x_i \Big\}}{1 - \exp\Big\{ - NA \langle d^{-1}\rangle \Big\}}. \qquad (64)
\end{aligned}$$

Assuming that the mutant was introduced in a random node of the network, the fixation probability can be written as

$$P\left(\vec{x} = \frac{\vec{1}}{N}\right) = \frac{1 - \exp\Big\{ - A\langle d^{-1}\rangle \Big\}}{1 - \exp\Big\{ - NA\langle d^{-1}\rangle \Big\}}. \qquad (65)$$

To summarize, the fixation probability for the Birth-death process on a network is given by

$$P_{Bd} = \frac{1 - e^{-\alpha_{Bd} s/(1+s/2)}}{1 - e^{-\alpha_{Bd} N s/(1+s/2)}}, \qquad \text{where } \alpha_{Bd} = \left(\langle d^{-1}\rangle \sum_{i,j\in D} p_j p_{ji} d_i^{-1}\right)\left(\sum_{i,j\in D} p_j p_{ji} d_i^{-2}\right)^{-1}. \qquad (66)$$

Here, $\alpha_{Bd}$ is the network quantity that governs the evolutionary dynamics on graphs under the Birth-death update rule.

In **Supplementary Figure S7** we show how well (66) approximates the fixation probability obtained from solving (53) numerically. In our derivation of the approximation, we ignored the $\mathcal{O}(s^2)$ portion of the roots of (53). The error that accumulates is on the order of $Ns^2$, therefore, as long as $s << N^{-1/2}$ the approximation should hold. The numerical solution starts to deviate from the approximate solution as $s$ increases for $\alpha_{Bd} < 1$.

#### Change in amplification due to rewiring

Here we expand on the derivation of equation (11) in the main text. We write the numerator and denominator in (66):

$$\mu_1 = \sum_{i,j \in D} p_j p_{ji} d_i^{-1},$$
$$\mu_2 = \sum_{i,j \in D} p_j p_{ji} d_i^{-2} \tag{67}$$

and we consider the change in the numerator and denominator under one rewiring step:

$$\begin{aligned}
\Delta\mu_1 &= -p_i \frac{1}{Np_i d_i}\frac{1}{d_i} - p_j \frac{1}{Np_j d_j}\frac{1}{d_j} + p_i \frac{1}{Np_i d_i}\frac{1}{d_j} + p_j \frac{1}{Np_j d_j}\frac{1}{d_i} \\
&= -\frac{1}{Nd_i^2} - \frac{1}{Nd_j^2} + \frac{2}{Nd_i d_j} \\
&= \frac{1}{N}\frac{2d_i d_j - d_i^2 - d_j^2}{d_i^2 d_j^2} \\
&= -\frac{1}{N}\frac{(d_i - d_j)^2}{d_i^2 d_j^2} < 0
\end{aligned}$$

and

$$\begin{aligned}
\Delta\mu_2 &= -p_i \frac{1}{Np_i d_i}\frac{1}{d_i^2} - p_j \frac{1}{Np_j d_j}\frac{1}{d_j^2} + p_i \frac{1}{Np_i d_i}\frac{1}{d_j^2} + p_j \frac{1}{Np_j d_j}\frac{1}{d_i^2} \\
&= -\frac{1}{Nd_i^3} - \frac{1}{Nd_j^3} + \frac{1}{Nd_i^2 d_j} + \frac{1}{Nd_i d_j^2} \\
&= \frac{1}{N}\frac{d_i^2 d_j + d_i d_j^2 - d_i^3 - d_j^3}{d_i^3 d_j^3} \\
&= \frac{1}{N}\frac{d_i^2(d_j - d_i) + d_j^2(d_i - d_j)}{d_i^3 d_j^3} \\
&= -\frac{1}{N}\frac{(d_i^2 - d_j^2)(d_i - d_j)}{d_i^3 d_j^3} \\
&= -\frac{1}{N}\frac{(d_i + d_j)(d_i - d_j)^2}{d_i^3 d_j^3} < 0.
\end{aligned}$$

Since the change is on the order of $\frac{1}{N}$, we can approximate the change by

$$\begin{aligned}
\Delta\frac{\mu_1}{\mu_2} &= \frac{\mu_1+\Delta\mu_1}{\mu_2+\Delta\mu_2} - \frac{\mu_1}{\mu_2} \\
&= (\mu_1+\Delta\mu_1)\left(\frac{1}{\mu_2} - \frac{\Delta\mu_2}{\mu_2^2}\right) - \frac{\mu_1}{\mu_2} \\
&= \frac{\Delta\mu_1}{\mu_2} - \frac{\mu_1\Delta\mu_2}{\mu_2^2} \\
&= \frac{\mu_1}{\mu_2^2}\left(\frac{\mu_2}{\mu_1}\Delta\mu_1 - \Delta\mu_2\right) \\
&= \frac{\mu_1}{N\mu_2^2}\left(\frac{\mu_2}{\mu_1}\frac{-(d_i-d_j)^2}{d_i^2 d_j^2} - \frac{(d_i+d_j)(d_i-d_j)^2}{d_i^3 d_j^3}\right) \\
&= \frac{\mu_1}{N\mu_2^2}\frac{(d_i-d_j)^2}{d_i^2 d_j^2}\left(-\frac{\mu_2}{\mu_1} + \frac{d_i+d_j}{d_i d_j}\right) \\
&= \frac{\mu_1}{N\mu_2^2}\frac{(d_i-d_j)^2}{d_i^2 d_j^2}\left(\frac{1}{d_i} + \frac{1}{d_j} - \frac{\mu_2}{\mu_1}\right).
\end{aligned}$$

### Approximation for detour graphs under weak selection

Here we present the derivation of equations (13) in the main text. We obtain an alternate approximate solution to the Kolmogorov backward equation by using regular perturbation. This is because the previous derivation underestimates probabilities of fixation on detour graphs, since they have very few edges that connect nodes of different degrees.

We expand the solution to (47) in terms of $s$

$$P = P_0 + sP_1 + s^2 P_2 + .... \tag{68}$$

Substitute into equation (47) and obtain the following

$$\sum_{i\in D, j\in D} p_j p_{ji}\left(\frac{1}{2Np_i^2}[(1+s)x_j+x_i-(2+s)x_i x_j]\frac{\partial^2}{\partial x_i^2}(P_0+sP_1+...)+\frac{1}{p_i}[(1+s)x_j-x_i-sx_i x_j]\frac{\partial}{\partial x_i}(P_0+sP_1+...)\right) = 0. \tag{69}$$

Under weak selection, the terms in the equation above independent of $s$ can be written as

$$\sum_{i\in D, j\in D} p_j p_{ji}\left(\frac{1}{2Np_i^2}(x_j + x_i - 2x_i x_j)\frac{\partial^2 P_0}{\partial x_i^2} + \frac{1}{p_i}(x_j - x_i)\frac{\partial P_0}{\partial x_i}\right) = 0. \tag{70}$$

This equation is identical to the Kolmogorov backward equation under neutrality. The solution is known and is given by

$$P_0 = \frac{1}{\langle d^{-1}\rangle} \sum_{i\in D} \frac{p_i x_i}{d_i}. \tag{71}$$

Next, we collect the terms of the first order term of $s$ and write

$$\begin{aligned}
&\sum_{i\in D, j\in D} p_j p_{ji} \Bigg( \frac{1}{2Np_i^2}(x_j - x_i x_j)\frac{\partial^2 P_0}{\partial x_i^2} + \frac{1}{p_i}(x_j - x_i x_j)\frac{\partial P_0}{\partial x_i} + \\
&\qquad \frac{1}{2Np_i^2}(x_j + x_i - 2x_i x_j)\frac{\partial^2 P_1}{\partial x_i^2} + \frac{1}{p_i}(x_j - x_i)\frac{\partial P_1}{\partial x_i} \Bigg) = 0 \\
&= \sum_{i\in D, j\in D} p_j p_{ji} \Bigg( \frac{1}{\langle d^{-1}\rangle d_i}(x_j - x_i x_j) + \\
&\qquad \frac{1}{2Np_i^2}(x_j + x_i - 2x_i x_j)\frac{\partial^2 P_1}{\partial x_i^2} + \frac{1}{p_i}(x_j - x_i)\frac{\partial P_1}{\partial x_i} \Bigg) = 0.
\end{aligned} \tag{72}$$

The solution has the form

$$\begin{aligned}
P_1 &= \sum_{ij} p_i p_j A_{ij} x_i (1 - x_j) \\
&= \sum_i p_i A_i x_i - \sum_{ij} p_i p_j A_{ij} x_i x_j, \text{ where } A_i = \sum_j p_j A_{ij}.
\end{aligned} \tag{73}$$

and we need to solve for the unknowns $A_i$ and $A_{ij}$. We know the solution to (72) has to have this form because the neutrality solution $P_0$ already satisfies the boundary conditions $P(0) = 0$ and $P(1) = 1$, so $P_1(0) = 0$ and $P_1(1) = 0$ are required. The partial derivatives are given by

$$\frac{\partial P_1}{\partial x_i} = p_i A_i - 2p_i \sum_j p_j A_{ij} \quad \text{and} \quad \frac{\partial^2 P_1}{\partial x_i x_j} = -2p_i p_j A_{ij}. \tag{74}$$

Substitute in (72) and we have

$$\begin{aligned}
&\sum_{i\in D, j\in D} p_j p_{ji} \Bigg[ \frac{1}{\langle d^{-1}\rangle d_i}(x_j - x_i x_j) - \frac{A_{ii}}{N}(x_j + x_i - 2x_i x_j) \\
&\qquad + (x_j - x_i)\Bigg( A_i - \sum_k p_k A_{ik} x_k - p_i A_{ii} x_i \Bigg) \Bigg] = 0.
\end{aligned} \tag{75}$$

In order for the equation to be satisfied all the coefficients must sum to zero. Therefore, the conditions for the linear terms, $x_i'$s, are

$$\sum_{j\in D}\left[-p_jp_{ji}\left(\frac{A_{ii}}{N}+A_i\right)+p_ip_{ij}\left(\frac{1}{\langle d^{-1}\rangle d_j}-\frac{A_{jj}}{N}+A_j\right)\right]=0. \tag{76}$$

For the quadratic terms, $x_ix_j'$s, we re-index to collect the like quadratic terms

$$\begin{aligned}
&\sum_{i\in D,j\in D}p_jp_{ji}\left(-\frac{x_ix_j}{\langle d^{-1}\rangle d_i}+2\frac{A_{ii}}{N}x_ix_j-2(x_j-x_i)\sum_k p_kA_{ik}x_k\right)=0\\
\implies&\sum_{i\in D,j\in D}p_jp_{ji}\left(-\frac{x_ix_j}{\langle d^{-1}\rangle d_i}+2\frac{A_{ii}}{N}x_ix_j-2\sum_k p_kA_{ik}x_jx_k+2\sum_k p_kA_{ik}x_ix_k\right)=0\\
\implies&\sum_{i\in D,j\in D}\Bigg[p_jp_{ji}\left(-\frac{x_ix_j}{\langle d^{-1}\rangle d_i}+2\frac{A_{ii}}{N}x_ix_j\right)\\
&\qquad-2\sum_k p_kp_jp_{ji}A_{ik}x_jx_k+2\sum_k p_kp_jp_{ji}A_{ik}x_ix_k\Bigg]=0\\
\implies&\sum_{i\in D,j\in D}\Bigg[p_jp_{ji}\left(-\frac{x_ix_j}{\langle d^{-1}\rangle d_i}+2\frac{A_{ii}}{N}x_ix_j\right)\\
&\qquad-2\sum_k p_ip_jp_{jk}A_{ki}x_ix_j+2\sum_k p_jp_kp_{ki}A_{ij}x_ix_j\Bigg]=0.
\end{aligned} \tag{77}$$

For the coefficients of the quadratic terms to sum to zero, the following set of equations must be satisfied

$$\begin{aligned}
&p_jp_{ji}\left(-\frac{1}{\langle d^{-1}\rangle d_i}+2\frac{A_{ii}}{N}\right)-2\sum_k p_ip_jp_{jk}A_{ki}+2\sum_k p_jp_kp_{ki}A_{ij}\\
&\qquad+p_ip_{ij}\left(-\frac{1}{\langle d^{-1}\rangle d_j}+2\frac{A_{jj}}{N}\right)-2\sum_k p_jp_ip_{ik}A_{kj}+2\sum_k p_ip_kp_{kj}A_{ij}=0.
\end{aligned} \tag{78}$$

Equations (74), (76), and (78) form a system of linear equations in which we solve for all the $A$ terms.

Next, we show that equation (76) is actually redundant given (74) and (78). To do so, we sum (78) by $j$

and apply (74) and write

$$\begin{aligned}
&\sum_j p_j p_{ji}\left(\frac{1}{\langle d^{-1}\rangle d_i} - 2\frac{A_{ii}}{N}\right) + \sum_j p_i p_{ij}\left(\frac{1}{\langle d^{-1}\rangle d_j} - 2\frac{A_{jj}}{N}\right)\\
&\qquad + 2\sum_{jk} p_i p_j p_{jk} A_{ki} + 2\sum_{jk} p_j p_i p_{ik} A_{kj} = 2\sum_{kj} p_i p_k p_{kj} A_{ij} + 2\sum_{kj} p_j p_k p_{ki} A_{ij}\\
&\implies \sum_j p_j p_{ji}\left(\frac{1}{\langle d^{-1}\rangle d_i} - 2\frac{A_{ii}}{N}\right) + \sum_j p_i p_{ij}\left(\frac{1}{\langle d^{-1}\rangle d_j} - 2\frac{A_{jj}}{N}\right)\\
&\qquad + 2\sum_{jk} p_i p_j p_{jk} A_{ki} + 2\sum_{k} p_i p_{ik} A_{k} = 2\sum_{kj} p_i p_k p_{kj} A_{ij} + 2\sum_{k} p_k p_{ki} A_{i}\\
&\implies \sum_j p_j p_{ji}\left(\frac{1}{\langle d^{-1}\rangle d_i} - 2\frac{A_{ii}}{N}\right) + \sum_j p_i p_{ij}\left(\frac{1}{\langle d^{-1}\rangle d_j} - 2\frac{A_{jj}}{N}\right)\\
&\qquad + 2\sum_{jk} p_i p_j p_{jk} A_{ki} + 2\sum_{k} p_i p_{ik} A_{k} = 2\sum_{kj} p_i p_j p_{jk} A_{ik} + 2\sum_{k} p_k p_{ki} A_{i}\\
&\implies \sum_j p_j p_{ji}\left(\frac{1}{\langle d^{-1}\rangle d_i} - 2\frac{A_{ii}}{N}\right) + \sum_j p_i p_{ij}\left(\frac{1}{\langle d^{-1}\rangle d_j} - 2\frac{A_{jj}}{N}\right) + 2\sum_k p_i p_{ik} A_k = 2\sum_k p_k p_{ki} A_i
\end{aligned} \tag{79}$$

Lastly, we set this equal to two times equations (76),

$$\begin{aligned}
&\sum_j p_j p_{ji}\left(\frac{1}{\langle d^{-1}\rangle d_i} - 2\frac{A_{ii}}{N}\right) + \sum_j p_i p_{ij}\left(\frac{1}{\langle d^{-1}\rangle d_j} - 2\frac{A_{jj}}{N}\right) + 2\sum_k p_i p_{ik} A_k - 2\sum_k p_k p_{ki} A_i\\
&\qquad = 2\sum_{j\in D}\left[ - p_j p_{ji}\left(\frac{A_{ii}}{N} + A_i\right) + p_i p_{ij}\left(\frac{1}{\langle d^{-1}\rangle d_j} - \frac{A_{jj}}{N} + A_j\right)\right] = 0\\
&\implies \sum_j p_j p_{ji}\left(\frac{1}{\langle d^{-1}\rangle d_i}\right) + \sum_j p_i p_{ij}\left(\frac{1}{\langle d^{-1}\rangle d_j}\right) + 2\sum_k p_i p_{ik} A_k - 2\sum_k p_k p_{ki} A_i\\
&\qquad = 2\sum_{j\in D}\left[ - p_j p_{ji} A_i + p_i p_{ij}\left(\frac{1}{\langle d^{-1}\rangle d_j} + A_j\right)\right]\\
&\implies \sum_j p_j p_{ji}\left(\frac{1}{\langle d^{-1}\rangle d_i}\right) + \sum_j p_i p_{ij}\left(\frac{1}{\langle d^{-1}\rangle d_j}\right) = 2\sum_{j\in D} p_i p_{ij}\left(\frac{1}{\langle d^{-1}\rangle d_j}\right)\\
&\implies \sum_j p_j p_{ji}\left(\frac{1}{\langle d^{-1}\rangle d_i}\right) = \sum_{j\in D} p_i p_{ij}\left(\frac{1}{\langle d^{-1}\rangle d_j}\right).
\end{aligned} \tag{80}$$

This last equation is true by the handshake lemma (see (29)). This proves that (76) is redundant given (74) and (78) and therefore we need only solve a much smaller set of equations.

To conclude, we can approximate the fixation probability using

$$P(x) \approx \frac{1}{\langle d^{-1} \rangle} \sum_{i \in D} \frac{p_i x_i}{d_i} + s \sum_{ij} p_i p_j A_{ij} x_i (1 - x_j), \tag{81}$$

where $A_{ij}$ is found by solving (78).

### Supplementary figures

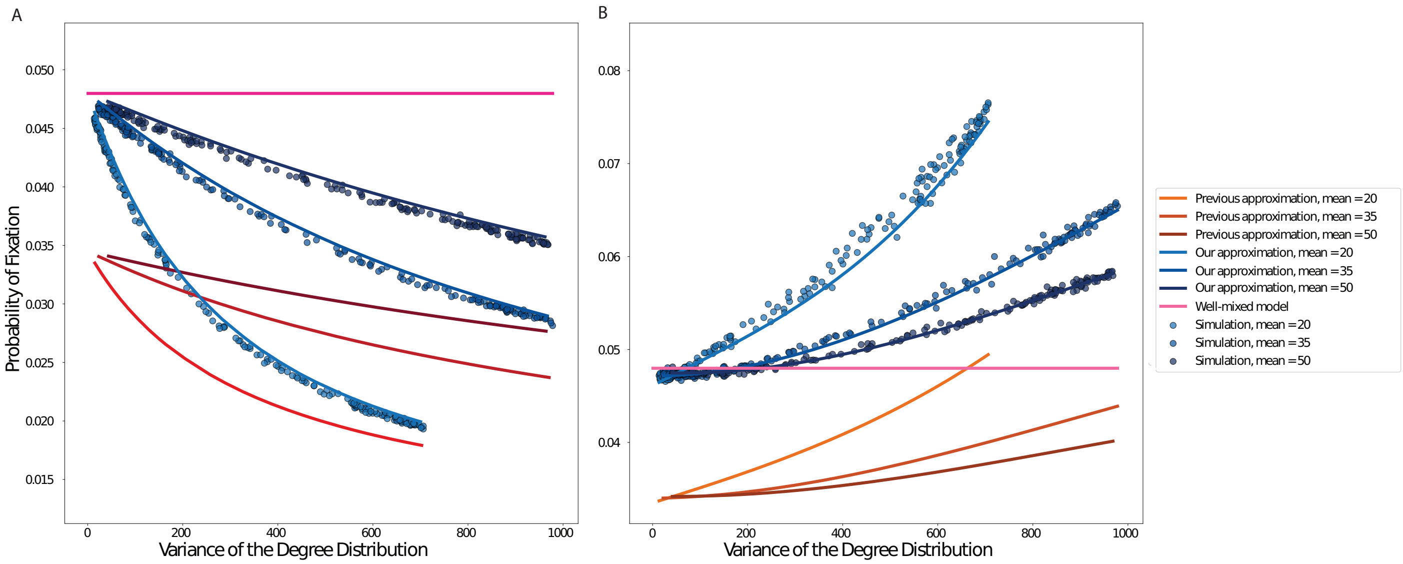


Figure S1: **Comparison with previous analytical methods.** The dots represent represent ensemble averages across $10^6$ replicate Monte Carlo simulations, while the lines represent our analytical approximations. **Panel A** corresponds to the death-Birth update rule, while **Panel B** shows results for thre Birth-death process. We use preferential attachment PA graphs, graph size $N = 100$ and $Ns = 5$.

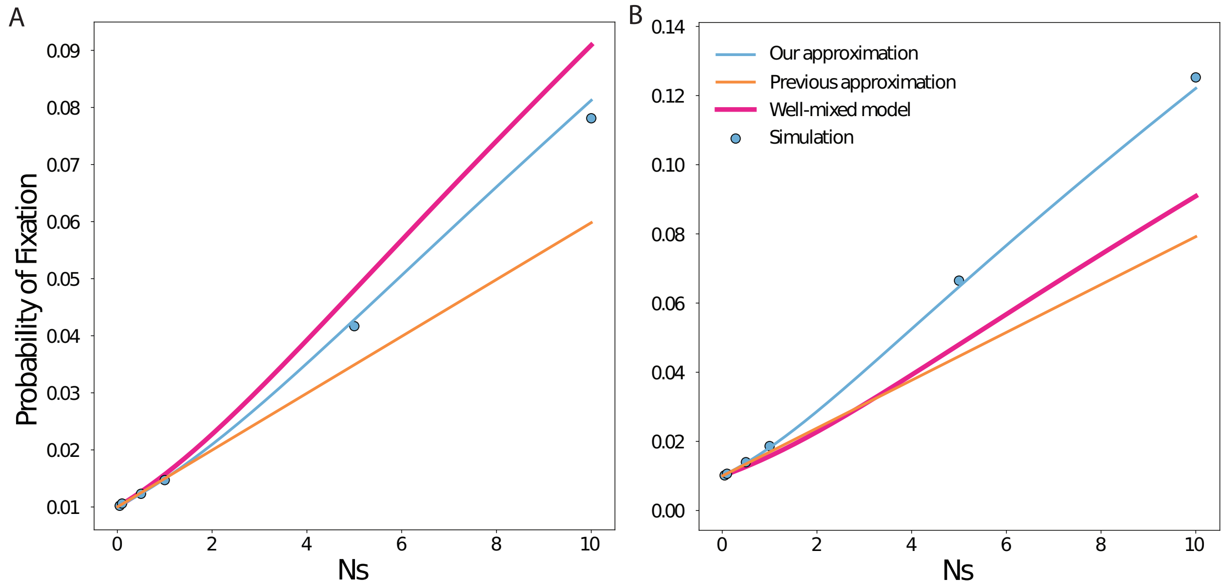


Figure S2: **Comparison with previous analytical methods.** The dots represent represent ensemble averages across $10^6$ replicate Monte Carlo simulations, while the lines represent our analytical approximations. **Panel A**: We show results for the death-Birth process on preferential attachment graphs with mean degree equal to 5.88 and variance in degree is 4.75. Graph size $N = 100$. $Ns$ ranges from 0.001 to 10. **Panel B**: We show results for the Birth-death process on preferential attachment graphs with mean degree equal to 5.88 and variance in degree is 266.3. Graph size $N = 100$. $Ns$ ranges from 0.001 to 10.

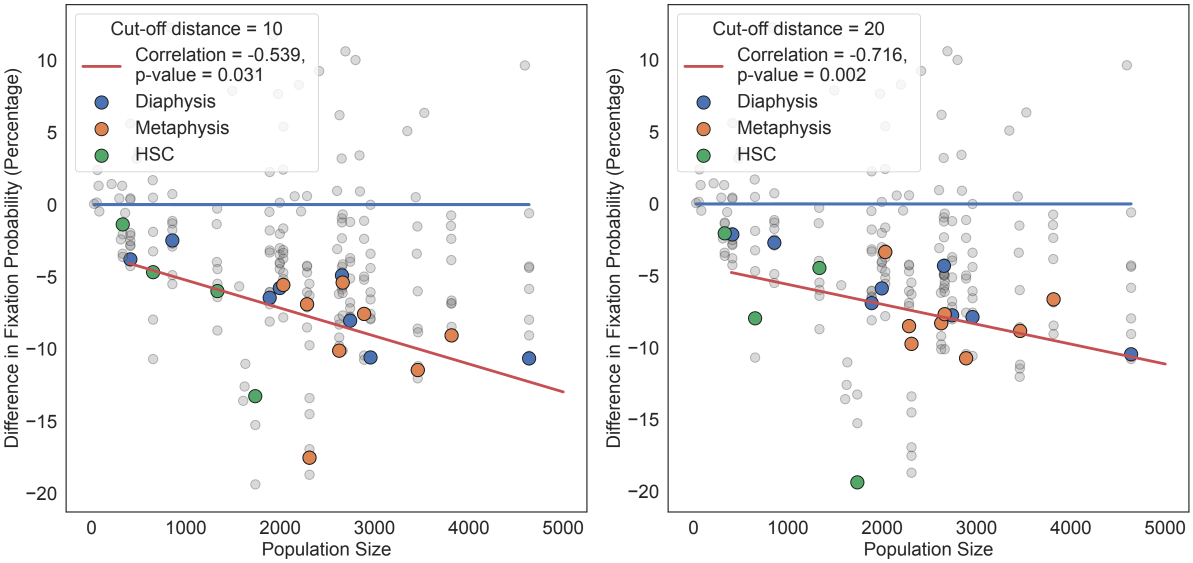


Figure S3: **Robustness of cutoff distance for the bone marrow networks.** Similar to Figure 6 in main text. Here we build the stem cell geometric random graphs and the color dots use cut-off distances of 10 and 20 for the two panels. Here, $s = 0.005$ and $Ns$ varies with population size.

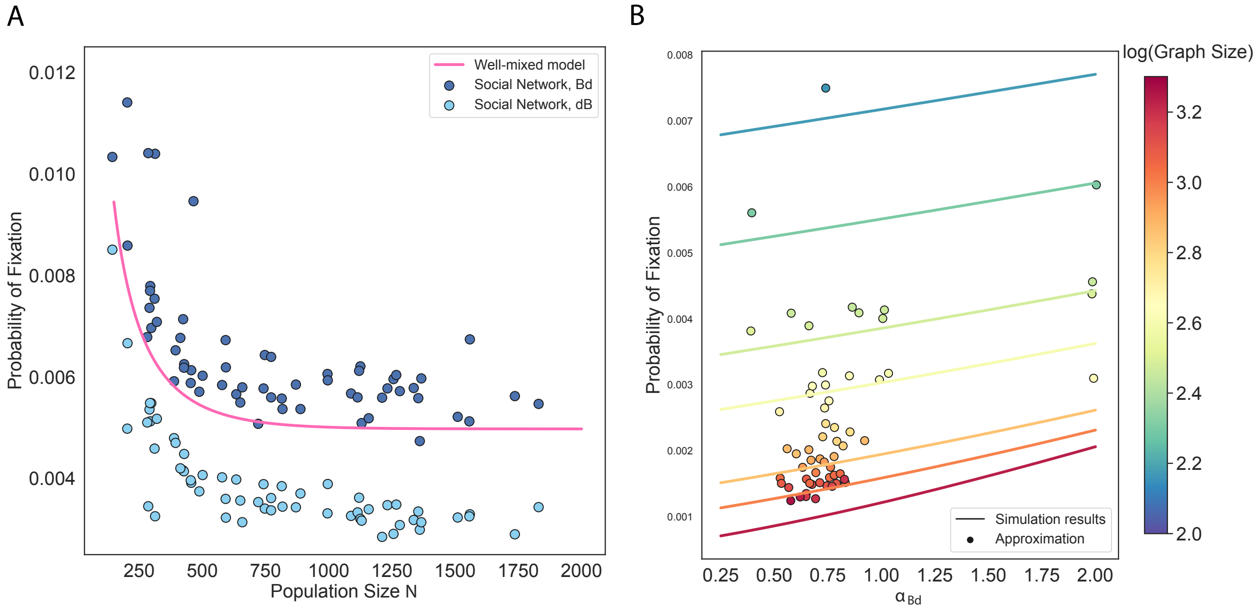


Figure S4: **Evolutionary dynamics on social networks.** Here we present results for social network datasets as described in the main paper Discussion. Fixation probability is calculated using a selective advantage of $s = 0.005$, while $Ns$ varies from 0.0625 to 5. The pink line is the probability of fixation in well-mixed populations. These networks are overwhelmingly amplifiers under the Birth-death process and suppressors under the death-Birth process.

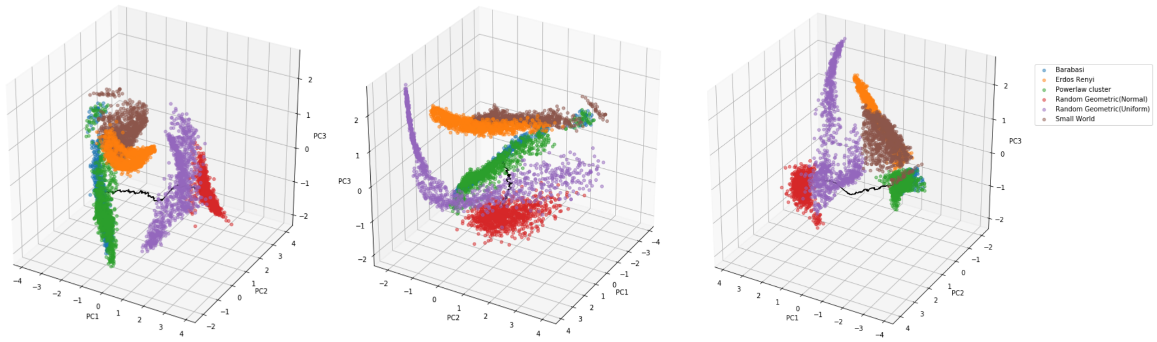


Figure S5: **Visualizing the space of network statistics explored.** We use principle component analysis on six graph characteristics (mean, variance, third moment, modularity, average clustering, and assortativity). Each graph family clusters together and we use novel network generation algorithms to explores the spaces in between generation algorithms that are family-specific. The black line represents a trajectory in PCA space of the rewiring from PA to RGG. The trajectory starts at PA and passes through PLC and RGG(uniform) to RGG(normal).

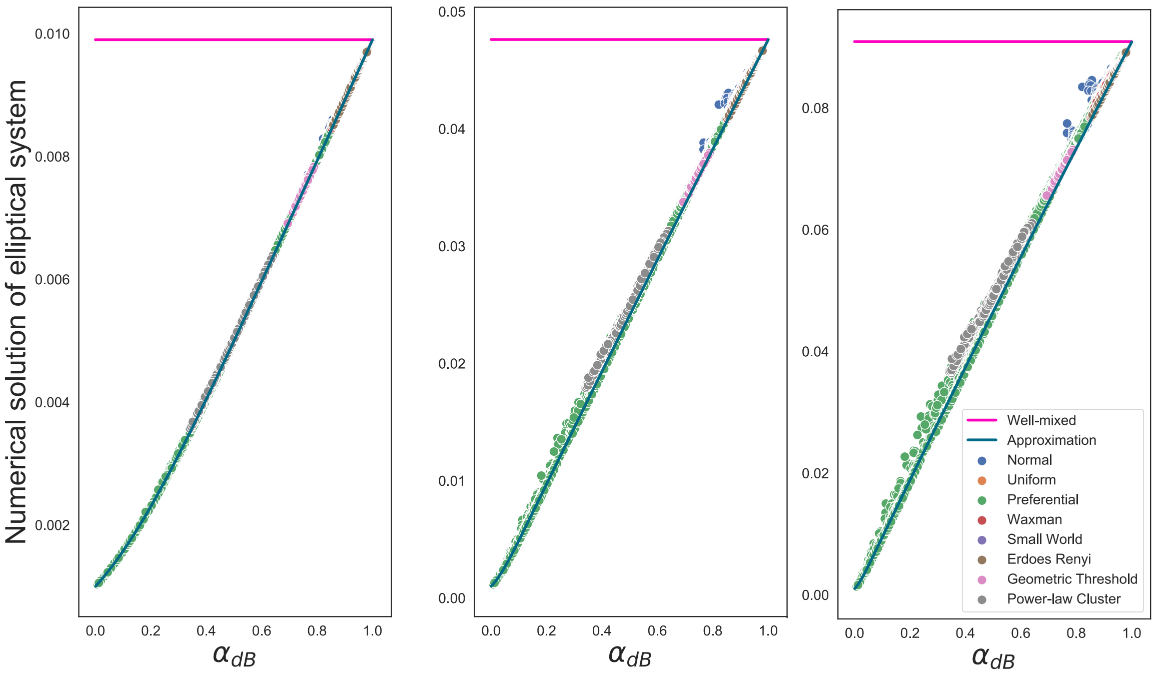


Figure S6: **Analytical approximation of the solution to the diffusion equation for the death-Birth process.** The lines are the approximation of fixation probabilities using (35). The dots are approximations using the numerical solutions of (21). Each dot represents a distinct graph. There are 5703 graphs presented. Graph size $N = 1000$. The various colors represent different network families. **Panel A** $s = 0.01$, $Ns = 10$; **Panel B** $s = 0.05$, $Ns = 50$; and **Panel C** $s = 0.1$, $Ns = 100$.

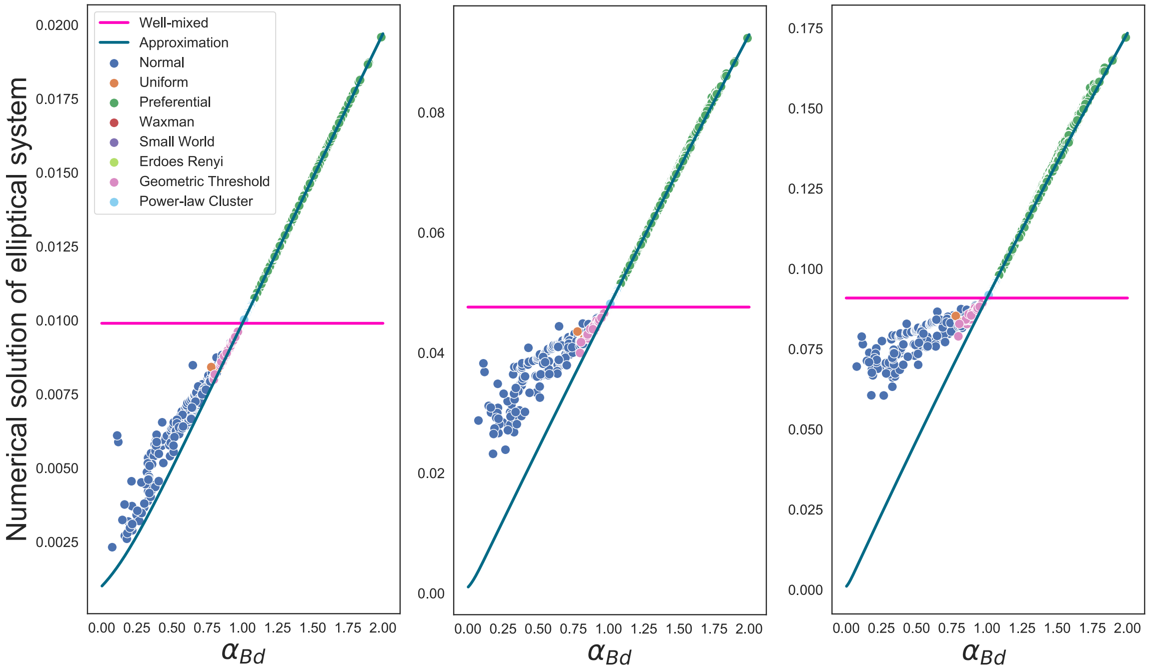


Figure S7: **Analytical approximation of the solution to the diffusion equation for the Birth-death process.** The lines are the approximation of fixation probabilities using (66). The dots are approximations using the numerical solutions of (53). Each dot represents a distinct graph. There are 5703 graphs presented. Graph size $N = 1000$. The various colors represent different network families. **Panel A** $s = 0.01$, $Ns = 10$; **Panel B** $s = 0.05$, $Ns = 50$; and **Panel C** $s = 0.1$, $Ns = 100$.

### List of Figures

Figure S1. **Comparison with previous analytical methods.** The dots represent represent ensemble averages across $10^6$ replicate Monte Carlo simulations, while the lines represent our analytical approximations. **Panel A** corresponds to the death-Birth update rule, while **Panel B** shows results for thre Birth-death process. We use preferential attachment PA graphs, graph size $N = 100$ and $Ns = 5$.

Figure S2. **Comparison with previous analytical methods.** The dots represent represent ensemble averages across $10^6$ replicate Monte Carlo simulations, while the lines represent our analytical approximations. **Panel A**: We show results for the death-Birth process on preferential attachment graphs with mean degree equal to 5.88 and variance in degree is 4.75. Graph size $N = 100$. $Ns$ ranges from 0.001 to 10. **Panel B**: We show results for the Birth-death process on preferential attachment graphs with mean degree equal to 5.88 and variance in degree is 266.3. Graph size $N = 100$. $Ns$ ranges from 0.001 to 10.

Figure S3. **Robustness of cutoff distance for the bone marrow networks.** Similar to Figure 6 in main text. Here we build the stem cell geometric random graphs and the color dots use cut-off distances of 10 and 20 for the two panels. Here, $s = 0.005$ and $Ns$ varies with population size.

Figure S4. **Evolutionary dynamics on social networks.** Here we present results for social network datasets as described in the main paper Discussion. Fixation probability is calculated using a selective advantage of $s = 0.005$, while $Ns$ varies from 0.0625 to 5. The pink line is the probability of fixation in well-mixed populations. These networks are overwhelmingly amplifiers under the Birth-death process and suppressors under the death-Birth process.

Figure S5. **Visualizing the space of network statistics explored.** We use principle component analysis on six graph characteristics (mean, variance, third moment, modularity, average clustering, and assortativity). Each graph family clusters together and we use novel network generation algorithms to explores the spaces in between generation algorithms that are family-specific. The black line represents a trajectory in PCA space of the rewiring from PA to RGG. The trajectory starts at PA and passes through PLC and RGG(uniform) to RGG(normal).

Figure S6. **Analytical approximation of the solution to the diffusion equation for the death-Birth process.** The lines are the approximation of fixation probabilities using (35). The dots are approximations using the numerical solutions of (21). Each dot represents a distinct graph. There are 5703 graphs presented. Graph size $N = 1000$. The various colors represent different network families. **Panel A** $s = 0.01$, $Ns = 10$; **Panel B** $s = 0.05$, $Ns = 50$; and **Panel C** $s = 0.1$, $Ns = 100$.

Figure S7. **Analytical approximation of the solution to the diffusion equation for the Birth-death process.** The lines are the approximation of fixation probabilities using (66). The dots are approximations using the numerical solutions of (53). Each dot represents a distinct graph. There are 5703 graphs presented. Graph size $N = 1000$. The various colors represent different network families. **Panel A** $s = 0.01$, $Ns = 10$; **Panel B** $s = 0.05$, $Ns = 50$; and **Panel C** $s = 0.1$, $Ns = 100$.
